# SMALL MOLECULES WHICH CAN KEEP THE ECTODOMAINS OF HUMAN FcγRIIIa “CLOSED”: THERAPY FOR AUTOIMMUNE THROMBOCYTOPENIA PURPURA?

**DOI:** 10.1101/2025.05.05.652134

**Authors:** Ashish

**Author notes:** Address correspondence to: Fnu Ashish, PhD Allo BioLabs LLC, Dallas TX USA 75204; www.allobiolabs.com.

## Abstract

Comparative analysis of crystal structures of the ectodomains of the unliganded Fcγ receptor A (FcγRIIIa), its complex to IgG Fc and two designer proteins named “affimers” revealed that if the two ectodomains of FcγRIIIa remain “closed”, then they are conformationally incompetent to bind Fc region of antigen or infected cell bound IgG and protect them from degradation in the endosomes. By tightly binding to its ectodomains, these affimers derailed binding of FcγRIIIa to Fc portion of IgG leading to latter’s degradation. This ability to decrease antibody titer suggests use of these affimers as therapeutic interventions in autoimmune conditions like immune thrombocytopenia purpura (ITP) which is caused by autoantibodies against platelets. I queried: are there any small molecules which can dock and lock the ectodomains of FcγRIIIa analogous to affimers? Applying MTiOpenScreen on few libraries, I shortlisted 30 drug-like molecules which preferentially docked in the cleft between ectodomains of human FcγRIIIa. Their complexes were subsequently analyzed by MD simulations at theoretical pH 7.4 (and 5.7) at 310 K. For all trajectories, we tracked two characteristic molecular descriptors of FcγRIIIa: θ and δ, the angle and distance between the center masses of the two ectodomains, respectively. The values of these descriptors, in conjunction with estimated binding energies from MD simulations allow me to report three new compounds which may act as and/or lead to small molecule interventions in autoimmune conditions like ITP.

**Graphical abstract:** 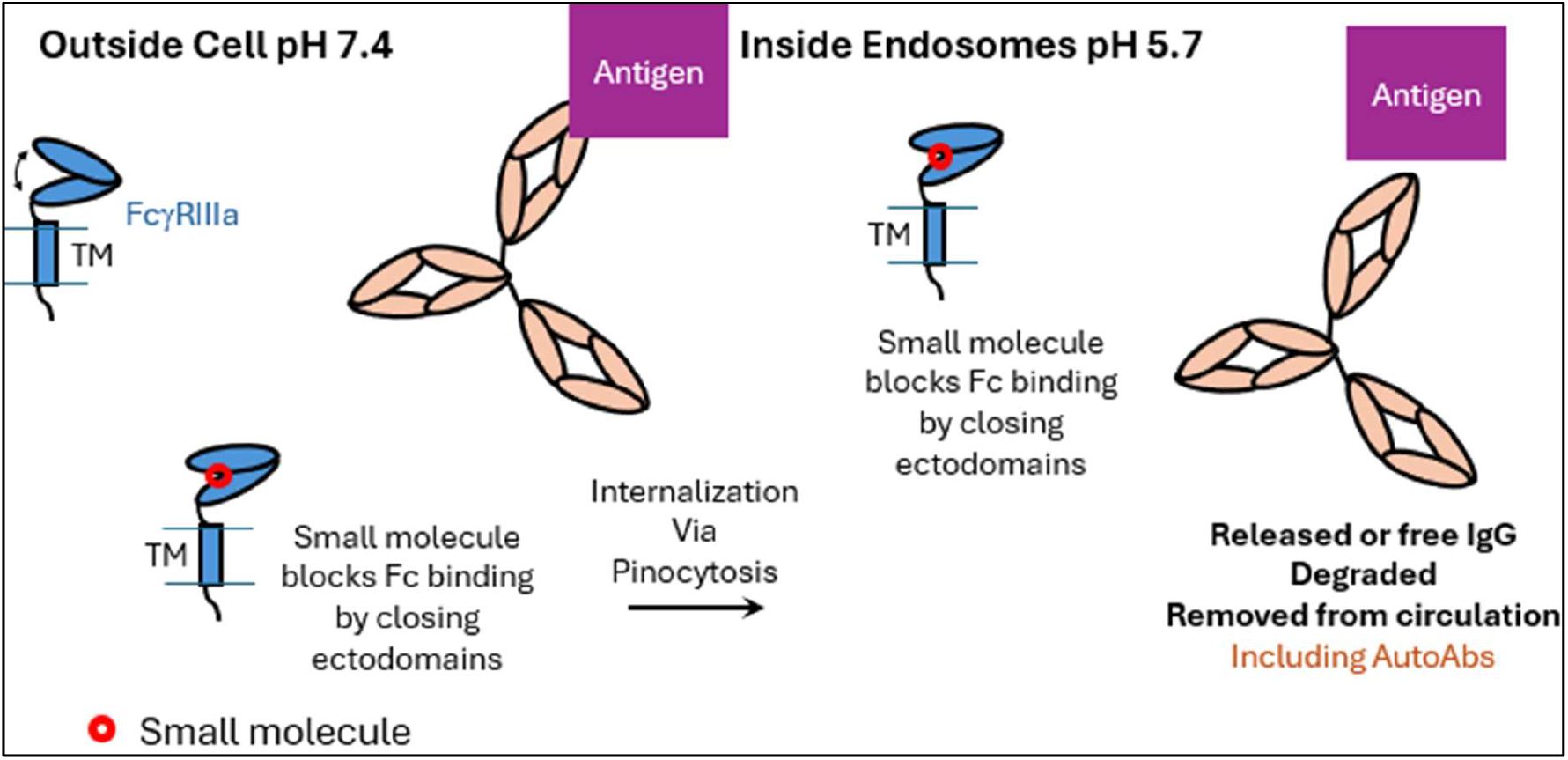

## Introduction

Fc gamma receptors (FcγRs) are surface proteins on immune cells that bind to different ligands or components on cells, triggering a variety of immune responses, including phagocytosis and antibody-dependent cellular cytotoxicity (ADCC) [1-3]. They primarily bind the Fc (fragment, crystallizable) region of IgG antibodies complexed to antigenic components on cells. Protein-protein interactions between FcγR and Fc of antigen-IgG complex are pivotal to many pro- and anti-inflammatory effects which arise via engagement of different FcγR [3]. In humans, there are six FcγRs which are sub-categorized into three main classes: FcγRI, FcγRIIa, FcγRIIb, FcγRIIc, FcγRIIIa, and FcγRIIIb [4]. During hominid evolution, different segmental deletions, duplications and post-translational modifications have resulted in families of highly homologous FcγR which can bind to different ligands and initiate varied biological functions [1-3,5]. This high degree of similarity in primary to tertiary structure has remained a problem while targeting specific FcγR for therapeutic interventions [1,6]. Earlier, a few monoclonal antibodies reactive to epitopes specific to FcγRs were tested for their ability to block antibody dependent cell cytotoxicity (ADCC) and treat ITP in humans [7]. Of those in antibody against FcγRIII, CD16-3G8 demonstrated early efficacy by increasing platelet counts. However, its usage was terminated due to observations of a typical hypersensitivity coupled with neutrophil and monocyte decrements. Even use of aglycosylated Fc portion to interfere with Fc-FcγRIII Interaction also showed similar results indicating that a better understanding is required in disrupting this pathway to develop an effective therapy for ITP [8]. More support came when a small molecule-based capable of blocking of spleen tyrosine kinase (SYK), a signaling molecule which occurs downstream of several FcγRs, showed some early promise in treating rheumatoid arthritis, chronic lymphocytic leukemia and non-Hodgkin’s lymphoma, yet further translation stopped due to adverse off-target results [9]. Despite failures, these studies hinted towards targeting interaction of Fc portions of IgG antibodies with FcγRIIIs to tackle with ITP.

FcγRIIIs have two highly homologous subsets, FcγRIIIa and FcγRIIIb, with about 98% sequence identity. While FcγRIIIa are expressed on the surfaces of macrophages and natural killer (NK) cells, FcγRIIIb are expressed on neutrophils. As can be seen in the sequence alignment shown in the **Supplementary Figure S1**, there are only 5 mutations across the 209 residues which compose the ectodomains of the two receptors. As mentioned earlier too, FcγRIIIa on macrophage surfaces bind to invariable Fc portion of the antigen-IgG complex and as a mega-complex anchored in membrane gets internalized into endosomes via pinocytosis where at low pH, lysis of antibody and antibody-coated cells is initiated, the process known as ADCC. It is important to mention here that FcγRIIIa does not bind free monomeric IgG, thus avoiding inappropriate degradation of antibodies or cell [10-12]. On NK cells, FcγRIIIa binds IgG-antigen complexes and triggers production of NK cell-dependent cytokine production [11,13]. In cases of viral infections, this pathway leads to production of IFNγ which enables elimination of virus infected cells via ADCC. It has been reported that during secondary infection with Dengue virus, elevated levels of afucosylated non-neutralizing IgG1 antibodies are produced which bind to FcγRIIIa with high affinity facilitating viral entry in myeloid cells and aiding in viral replication [14]. This antibody-dependent enhancement (ADE) of Dengue infection could be one reason for observing ITP cases post-Dengue infection. Thus, therapeutics and vaccines restricting production of afucosylated IgG1 and/or their binding to FcγRIIIa may prevent virus induced ADE [14]. In the **Supplementary Figure S2**, few schematic representations have been shown which summarize different approaches to block interaction between FcγRIIIa and antigen bound IgG Fc portion which will remove antibodies from circulation including autoantibodies raised in ITP. However, the big challenge is how to target FcγRIIIa specifically and not cross-react with FcγRIIIb?

In the inspirational work leading to this effort, the designer high affinity protein or affimers library, expressed on phages were screened against FcγRIIIa expressed on HEK 293T cells to search for high affinity binders specific to the ectodomains of this receptor [6]. Two affimers, AfG3 and AfF4 were identified to be the best binders, and their strong sub-micromolar scale binding affinities to FcγRIIIa were confirmed by isothermal titration calorimetry (ITC) and other affinity measurements. Importantly, by surface plasmon resonance assays (SPR), glycosylation on the ectodomains of the FcγRIIIa used for experiments did not confer significant difference in the binding profiles of the selected affimers. In other words, these affimers did not interact with epitopes proximal to glycosylation sites or sugar moieties in the ectodomains of FcγRIIIa. Further experiments showed that both AfG3 and AfF4 blocked binding of heat aggregated IgG1 (HAG) to FcγRIIIa, but not to FcγRIIIb or FcγRIIa expressed stably on HEK293 cells. The observed specificity could be visualized in the crystal structures of FcγRIIIa with AfG3 and AfF4, both in 1:1 ratio (PDB IDs 5MN2 and 5ML9, respectively) [6]. Analysis of the interaction sites further confirmed that they were specifically involving residues present in the ectodomains of FcγRIIIa, and absent in FcγRIIIb. Superimposition of the chains of FcγRIIIa in the crystal structures of FcγRIIIa bound to affimers AfG3, AfF4 and Fc portion of IgG, i.e. 5MN2, 5ML9 and 7URU, respectively showed that the binding pose of AfF4 on FcγRIIIa interferes with that of Fc portion on receptor (**Supplementary Figure S3 and Figure 1**). Overlap of the interacting epitopes and resultant steric interference explained the inhibition of Fc binding to FcγRIIIa by affimer AfF4. In sharp contrast, the affimer AfG3 binds on the opposite side of the Fc binding surface on FcγRIIIa, between its two ectodomains (PDB ID 5MN2) [6]. AfG3 has no direct interference with the Fc binding on FcγRIIIa. Instead, this affimer inhibits Fc binding allosterically. Analysis revealed that if the two ectodomains of FcγRIIIa are clasped together as induced by AfG3, their opposite surface involved in Fc binding is somehow rendered incompetent. Authors reported that the ability of FcγRIIIa to bind Fc portion of IgG1 can be correlated with the interdomain angle of FcγRIIIa [6]. They compared the opening between the two ectodomains of FcγRIIIa by measuring the angle between C^α^ atoms of Q83, W90 and N169 In the different crystal structures of unliganded FcγRIIIa, and it’s complex with IgG Fc and affimer AfG3. Authors found that in the crystal structures, interdomain angle was 46° in unliganded FcγRIIIa, which opened significantly to 53° in complex with Fc portion, and was substantially reduced to only 41° when bound to affimer AfG3. Taking together the structural data and other experimental results, it was a great finding that if the two ectodomains of FcγRIIIa can be held together, it may derail downstream events leading to ITP. With this very thought in mind, this *in silico* work was carried out to search for small drug like molecules which may dock between the two ectodomains of FcγRIIIa and hold them together?

**Figure 1.**
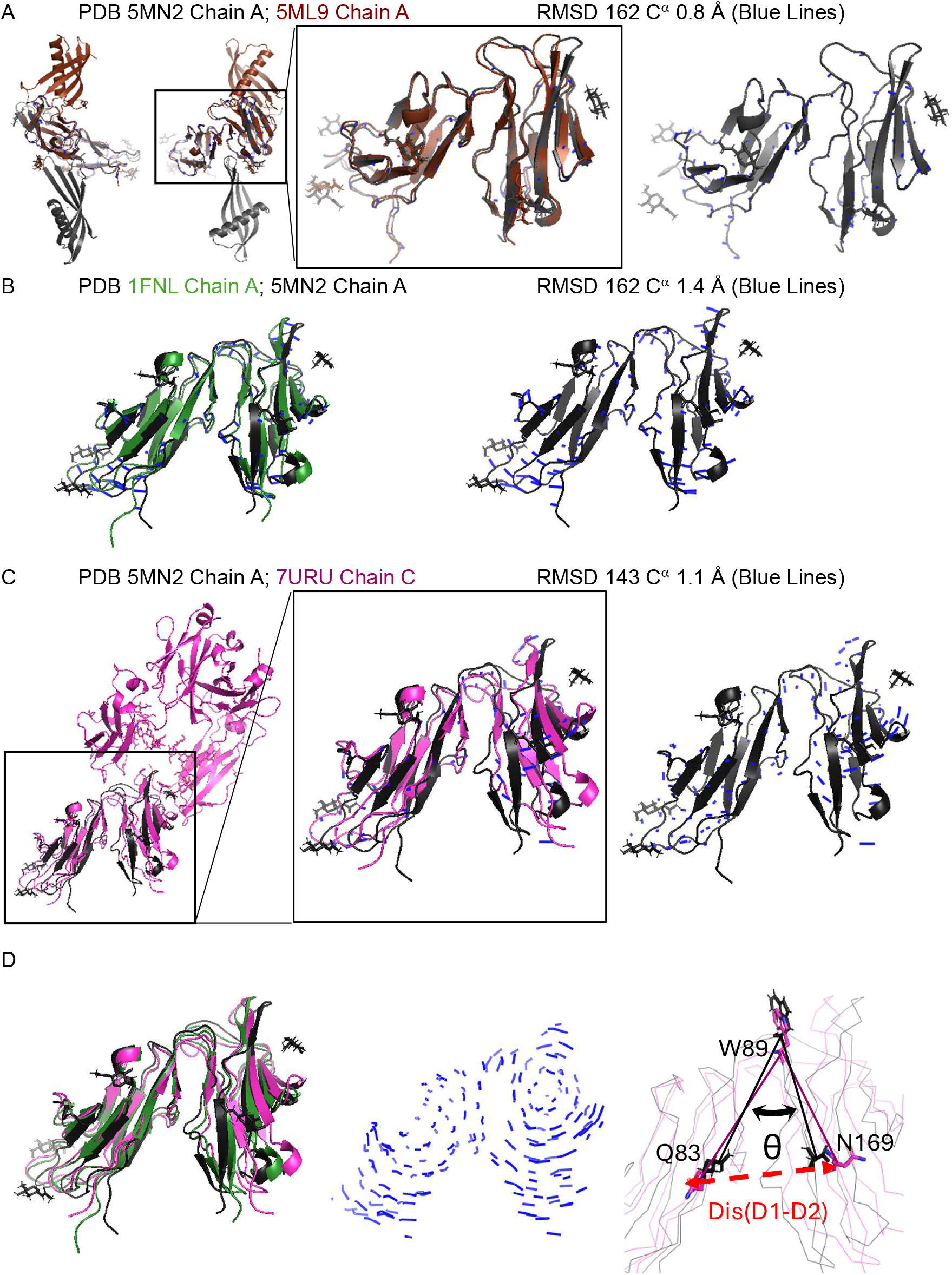
Crystal structures of Ectodomains of Human FcγRIIIa are compared here. (A) Structures of FcγRIIIa solved in complex with two high affinity affimers are shown here. Their PDB IDs are mentioned above in color of the structure. The left panel shows two rotated views of the complexes, and the box outlines the coordinates of the ectodomains of FcγRIIIa. The box in the middle panel shows the superimposition of the receptor in the two structures. The calculated RMSD values between the C^α^ residues are mentioned above, and the deviations are shown as blue lines with receptor from PDB ID 5MN2 as reference. (B) Structure of unglycosylated FcγRIIIa (PBD ID 1FNL) is compared with the glycosylated FcγRIIIa from PDB ID 5MN2 in left panel. The right panel shows the backbone deviations between the protein structures. (C) Left panel shows the superimposition of the FcγRIIIa portion of the complex with Fc with receptor from PDB ID 5MN2 (highlighted with box). As in part A, superimpositions and deviations are shown in reference to receptor from PDB ID 5MN2. (D) FcγRIIIa structures are superimposed to show deviations which are shown as blue lines in the central panel. The right panel shows the two descriptors which gives a measure of the closed structure of the FcγRIIIa ectodomains: θ, angle between Cα atoms of residues of Gln83, Trp89 and Gln169, and δ, the distance between the center of masses of domains D1 and D2.

## Methods

### Molecular Dynamics (MD) Simulations of FcγRIIIa -/+ Fc or Affimers

Different crystal structures were downloaded from the PDB database to extract structural coordinates of the ectodomains of FcγRIIIa. They were PDB ID 1FNL Chain A, 7URU Chain C, 5MN2 Chain A and 5ML9 Chain A representing unglycosylated FcγRIIIa, and glycosylated structures of FcγRIIIa bound to Fc portion of IgG and affimers AfG3 and AfF4, respectively. These structures were cleaned for any missing residues, bad contacts and all hydrogens were added to the structures. Additionally, PDB structures of ectodomains of FcγRIIIa complexed to Fc portion and the two affimers were also saved retaining the contacts in their crystal structures and cleaned. All energy minimizations, MD simulations and trajectory analysis were done using YASARA program version 24.4.10.W.64 [15,16] on a multi core and GPU assisted system [17,18]. All cleaned models of FcγRIIIa -/+ their binding partner were used at starting structures individually for a simulation time of 20 nanoseconds each using YAMBER2 force field at 310 K. The glycosyl moieties present in the crystal structures of FcγRIIIa were retained during simulations. All-atoms were considered in TIP3P water with pH set to 7.4 (and 5.7). NaCl concentration to 0.9% with additional monovalent ions were used to neutralize the overall charge of the system. Simulation cells were generated with target protein or complex in center and a cuboid geometry with extensions of 10 Å in each axis. Pressure on cell was regulated to maintain a cell density of ∼0.997 g/cc. Controls were enabled to reduce FcγRIIIa or its complex from drifting around and crossing periodic boundaries. Coordinates were integrated at timesteps of 2.5 femtoseconds. Calculation of non-bonded interactions were cut-off at 8 Å and periodic boundary conditions were considered to mimic simulation cell surroundings. After simulations and skipping the first 20 frames of saved snapshots, basic analysis was performed using macro-based routines. Different structural parameters like potential energy, radius of gyration (R_g_) of the solute, H-bonding profiles, RMSF values, secondary structural contents, nature of interactions, dynamic cross-correlation matrix etc. were calculated. Additionally, two characteristic molecular descriptors of FcγRIIIa: θ and δ, the angle and distance between the center masses of the two ectodomains, respectively. For θ, the angle between the C^α^ atoms of Q83 in D1 domain, W90 (fulcrum) and N169 in D2 domain of FcγRIIIa was measured for each snapshot. Similarly, distance between the center of masses of D1 domain (residues L5 to H87) and D2 domain (residues W90 to Q174) were measured to get value of δ. Later, the computed values of these molecular descriptors were plotted as a function of the simulation time using SAS plot software v 3.7.2.

### Docking Experiments

All docking experiments described in this work were done using MTiOpenScreen webserver [19]. For target, the average structure of glycosylated FcγRIIIa from MD simulation of the abstracted receptor from the PDB ID 5MN2 was considered. To focus the docking between the two ectodomains of FcγRIIIa, i.e. the epitope between the cleft of the two domains, surface formed by Q83, W90 and N169 was considered in a biased manner. Ligands with lead-like properties were prioritized and rotations about rotatable bonds were allowed. Three sets of compounds or libraries: drug-like, diverse compounds and food source were considered. 7173, 11000 and 10874 ligands from these respective libraries were considered for interaction with the epitope defined in the structure of FcγRIIIa. Results were provided in a formatted manner based on interaction energy and lowest energy docks were considered. In case multiple poses were provided for the same ligand, then the lowest energy pose was considered for further analysis. Finally, the top 10 ligands from each library were selected for MD simulations and subsequent analysis.

### MD Simulations of FcγRIIIa and selected ligands

All 30 complexes of glycosylated FcγRIIIa, each with one ligand docked in between the two ectodomains were cleaned, hydrogens were added, parameterized using YAMBER2 forcefield and energy minimized. Using the same macros as before, MD simulations were run for all the complexes and downstream processing were done. Additionally, binding constant estimations of each ligand to the FcγRIIIa receptor was done using the snapshots of its MD simulation trajectory. Few selected complexes of ligands to FcγRIIIa were re-simulated under pH conditions of 5.7.

### MD simulations of FcγRIIIb -/+ binding partners

The structure of the FcγRIIIb receptor was abstracted from PDB ID 6EAQ [20]. The structure of the two ectodomains of FcγRIIIb were solved bound to Fc portion with glycosylation in both the proteins. There were about eight residues missing in the D1 domain of FcγRIIIb. Using the crystal structure of the ectodomains as template and FASTA sequence of human FcγRIIIb (**Supplementary Figure S1**), SWISS MODELLER webserver was used to generate a model of FcγRIIIb without breaks in its 3D structure [21,22]. This structure was superimposed over the FcγRIIIb chain in the PDB ID 6EAQ to generate model of glycosylated FcγRIIIb, and latter’s complex with its Fc portion. All three structures, unglycosylated and glycosylated FcγRIIIb, and complex of glycosylated FcγRIIIb with Fc portion were subjected to MD simulations as described earlier at 310 K and pH of 7.4. Also, complex of glycosylated FcγRIIIb generated by retaining docking pose of selected ligands as on FcγRIIIa were analyzed by MD simulations and follow-up analysis. The two molecular descriptors, θ and δ were calculated for each snapshot in the trajectories but now using respective residues in the sequence of FcγRIIIb.

## Results and Discussion

### Ectodomains of FcγRIIIa from crystal structures and MD simulations at pH 7.4

Different crystal structures of FcγRIIIa as complex or unliganded are shown in cartoon mode in **Figure 1**. The two crystal structures of FcγRIIIa complexed to affimers, AfG3 and AfF4 i.e. PDB IDs 5MN2 and 5ML9, respectively are shown in **Figure 1A**. 3D structural alignment of the FcγRIIIa chains in the two structures showed an RMSD value of only 0.8 Å Indicating high degree of similarity in the structure of the ectodomains of FcγRIIIa despite bound to different affimers. Then, we compared the structure of FcγRIIIa in their aglycosylated and glycosylated form by aligning PDB IDs 1FNL and 5MN2, respectively (**Figure 1B**). A computed RMSD value of 1.4 Å indicated that the structure of the protein part of FcγRIIIa was largely same with or without glycosylation. Similarly, alignment of the ectodomains of FcγRIIIa bound to Fc portion of IgG and affimer AfG3 from the PDB IDs of 7URU and 5MN2, respectively computed an RMSD of only 1.1 Å which supported almost similar structure of FcγRIIIa between the two crystal structures (**Figure 1C**). In all the pairwise alignments, the spatial deviations between the positions of respective two C^α^ atoms are shown as blue lines. They imply that the deviations are relatively higher at the terminal ends and less around the linker portion of the two ectodomains. All deviations are shown together in the **Figure 1D** central panel which reflects the extent of breathing in the crystal structures of FcγRIIIa influenced by binding partner. It is pertinent to mention here that the observed RMSD values are average of the deviations in the pair of structures being aligned, hence a low RMSD value indicates similarity of the overall structure. It is important to mention here that the deviations in opposite directions may nullify and decrease the final summation. To obtain a better insight into the conformational preferences of FcγRIIIa as induced by its binding partner, we opted to measure two molecular descriptors, θ and δ, during their MD simulation trajectories, where they could be studied devoid of lattice packing effects known to influence crystal structures.

Results from the MD simulations of FcγRIIIa -/+ Fc portion or affimers done at 310 K and pH 7.4 and 5.7 are shown in **Figure 2**. The variations in the calculated values of molecular descriptors as a function of simulation time can be seen in the different panels. While the interdomain angle, θ in the trajectories started with the structures of FcγRIIIa extracted from its unliganded and affimers bound crystal structures wobbled close to 44-45° at pH 7.4, the structure extracted from the Fc bound crystal structure showed significantly higher values around 52° (**Figure 2 Top left**). Around simulation time of 10-15 ns, the θ values seem to be closer to 50 degrees for trajectories started with FcγRIIIa from crystal structures bound to AfG3 or Fc portion. Later, the θ values decreased for the simulation started using FcγRIIIa from PDB ID 5MN2, but the one initiated with FcγRIIIa from PDB ID 7URU indicated higher values or gap between the two ectodomains. Results indicated that there was significant amount of flexibility about the linker connecting the two ectodomains of FcγRIIIa. Our simulations indicated that glycosylation moieties around the protein part of FcγRIIIa did not influence significantly the interdomain angle which correlated with experimental binding studies [6]. Interestingly, the θ values were starkly different when the simulations were done considering the complexes of FcγRIIIa as refined in their PDB structures (**Figure 2 Top right**). The Fc portion kept the interdomain angle, θ substantially open close to 56°, bound affimers induced the ectodomains to remain closed which was reflected in θ values close to 41-43°, regardless of their binding pose on FcγRIIIa. The other molecular descriptor which we tracked was δ, the distance between the center of masses of the two ectodomains of FcγRIIIa. The trends correlated with the interdomain angle as there was higher variation in the δ values when the receptor was simulated after abstraction from the crystal structure(s) versus simulations carried out of the complexes. The main difference was that the δ values remained close to 24-25 Å in simulations carried out with unliganded receptor or as complex to affimers, but the structure solved with Fc portion showed slightly higher values. As before, interdomain movement was much less when complexes were simulated and observed influence of the two affimers were similar regardless of their binding pose on the FcγRIIIa.

**Figure 2.**
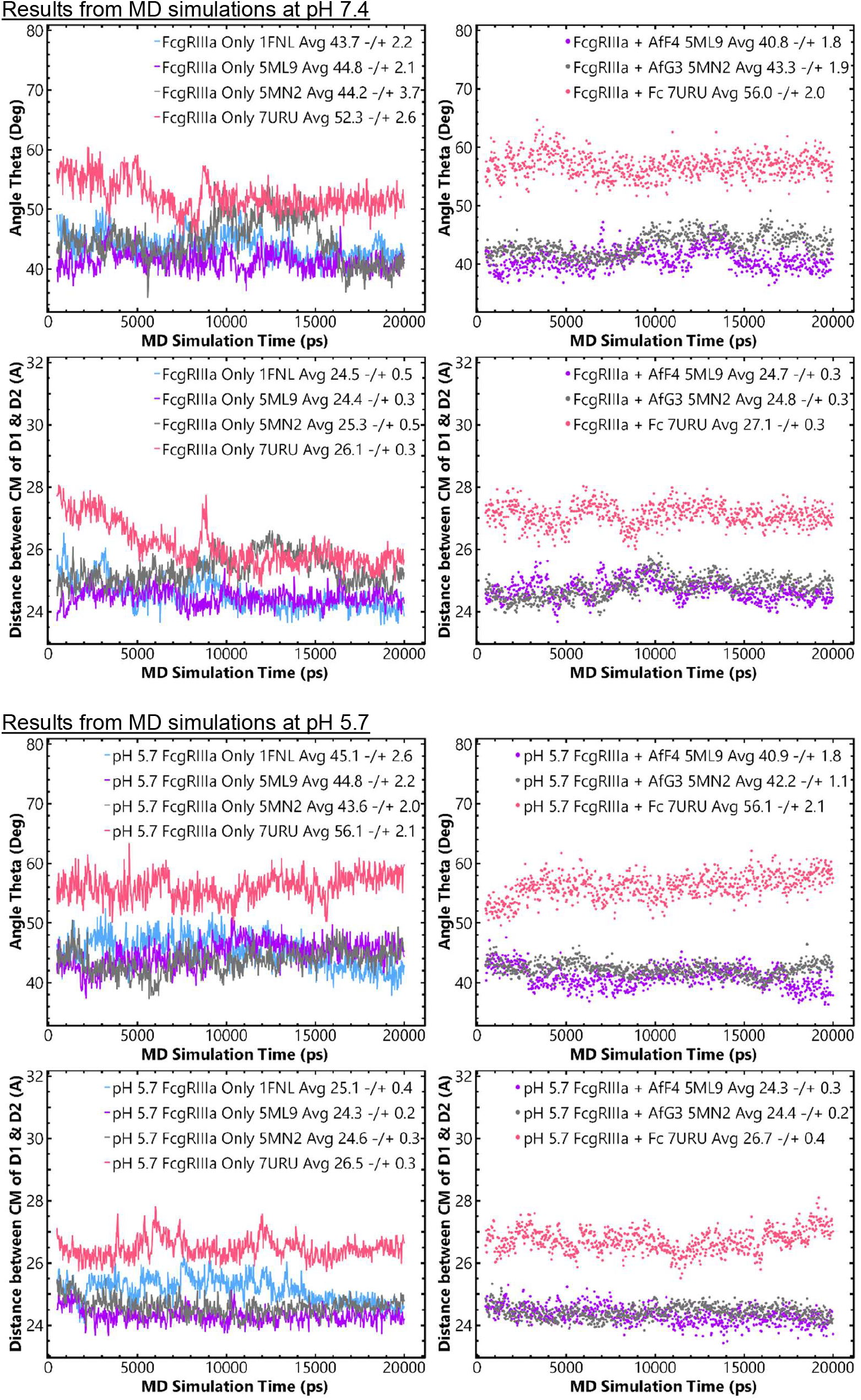
Variation in the calculated values of θ and δ in the simulated structures of the FcγRIIIa ectodomains as a function of the MD simulations at pH 7.4 and 5.7 are plotted here. Upper and lower four panels show variations in snapshots simulated at pH 7.4 and 5.7, respectively. Left and right panels show the values of the molecular descriptors for the unliganded receptor and their complexes, respectively. In the legends of the plots, the start structure of the calculation, average and standard deviation of the calculated descriptor are mentioned.

Since, the membrane anchored FcγRIIIa gets internalized and encounters pH close to 5.7 in the endosomes, to understand influence of lower pH on conformations of FcγRIIIa -/+ binding partners, additional MD simulations were done at lower pH, and the molecular descriptors were tracked (**Figure 2 Lower panels**). Primary observations were similar for both the descriptors, θ and δ at both the simulated pH values. Between the two pH sets, the main difference was the interdomain angle and distance did not reduce much for the FcγRIIIa structure abstracted from the crystal structure 7URU (**Figure 2 Lower panels**). Despite the differences in the variation patterns of the angle and distance between the two domains of unbound FcγRIIIa, at pH 5.7 (vs. pH 7.4), the θ values remained in the range of 43 to 45° and the δ values were close to 24 to 25 Å. As before, the movement between the two ectodomains of FcγRIIIa was substantially reduced in complexed states. There was little difference In the values of molecular descriptors in the two complexes of FcγRIIIa and affimers regardless of their binding preference on receptor and the bound Fc kept the domains of FcγRIIIa significantly open. The calculated R_g_ values of the FcγRIIIa structures as abstracted and as complexed to their binding partners are plotted in **Supplementary Figure S4**. The profiles and average values suggest that the R_g_ values were not significantly different in the receptor alone simulations and in the complexes. As expected, they reflect the bigger size of the complex with Fc portion vs. affimers. In other words, as seen for RMSD values of FcγRIIIa receptor in crystal structures, the R_g_ was not a good reflection of conformational changes in comparison to the molecular descriptors mentioned above. Importantly, comparison of the above four panels in the **Figure 2** indicated that the inherent flexibility of the FcγRIIIa receptor gets subdued, significantly and specifically by its binding partner, and those influences can be seen in the simulation times done here.

### Docking of ligands on FcγRIIIa and MD Simulations

As described in the Methods section, three libraries were docked inside the acute angle side of the ectodomains of FcγRIIIa. As target, the average structure of the partly glycosylated FcγRIIIa from the MD simulation of the receptor abstracted from PDB ID 5MN2 was used. From the results generated for each library 10 top poses of the docked ligands were selected for further analysis via MD simulations of the complexes. The shortlisted molecules from different libraries are tabulated in the **Supplementary Table ST1**. Their given names for this work are also mentioned in the tables. ZINC database IDs are mentioned for the available ones, and for the diverse molecule library hits, MTiOpenScreen IDs are mentioned. The final pose of the selected ligand(s) along with the target receptor structure was used as input files for MD simulations at pH 7.4. Parameterization was done for each ligand as docked/complexed to FcγRIIIa structure using YAMBER2 force field. Using macros, the molecular descriptors, θ and δ were calculated for each complex and are mentioned in **Table 1**. Receptor structure abstracted from the PDB ID 5MN2 showed average values of θ and δ 44.2° and 25.3 Å, respectively. Since our working hypothesis is that molecules which can close the interdomain gap of FcγRIIIa might interfere in the protection of autoantibodies and thus provide relief from ITP, we focused on molecules which can keep the two molecular descriptors of FcγRIIIa below 44.2° and 25.3 Å. Interestingly, a number of shortlisted ligands showed that during MD simulations they could keep the two ectodomains closer than seen for the unliganded FcγRIIIa. At this point, we added another criterion, i.e. the average binding energy of the snapshots from MD simulations (**Table 1**). Considering the cutoff average binding energy to be -200 kJ/mol, D4, V4 and V7 ligands were shortlisted as lead candidates. At this point, it is important to mention that none of the molecules selected from docking the Food library fitted over selection criteria. Rather in some cases, the two molecular descriptors increased in their values instead of expected decreasing. Whether these compounds or similar moieties can (artificially) cause aberrant behavior in signaling via FcγRIIIa pathways will remain a thought to ponder upon.

**Table 1.** Molecular descriptors of the structure of FcγRIIIa as complexed to their docked ligands are tabulated below. Also mentioned are the calculated average binding energy of the ligand to the receptor during MD simulation. Rows of the selected ligands are highlighted in grey shade.

| Drug-like Compounds | Angle $\theta$ (°) | $\delta$<br>CM D1D2 (Å) | MDS Based<br>Average Binding<br>Energy (kJ/mol) | Rating |
| --- | --- | --- | --- | --- |
| *Simulations done at pH 7.4* |  |  |  |  |
| **Approved Drug Library** |  |  |  |  |
| D1 | 44.5 ± 1.7 | 25.4 ± 0.3 | -177.26 |  |
| D2 | 43.5 ± 1.4 | 24.5 ± 0.2 | -189.32 | + |
| D3 | 43.7 ± 1.4 | 24.8 ± 0.2 | -182.10 | + |
| D4 | 41.3 ± 1.5 | 24.8 ± 0.3 | -214.70 | ++ |
| D5 | 46.3 ± 2.2 | 25.2 ± 0.3 | -154.25 |  |
| D6 | 44.8 ± 3.0 | 25.2 ± 0.3 | -142.36 |  |
| D7 | 43.2 ± 1.6 | 24.4 ± 0.4 | -190.81 | + |
| D8 | 47.4 ± 1.6 | 24.7 ± 0.2 | -085.05 |  |
| D9 | 45.8 ± 1.8 | 25.1 ± 0.2 | -111.75 |  |
| D10 | 45.6 ± 1.3 | 25.2 ± 0.3 | -109.21 |  |
| **Diverse Library** |  |  |  |  |
| V1 | 43.2 ± 1.9 | 24.9 ± 0.3 | -180.11 | + |
| V2 | 46.0 ± 2.0 | 25.0 ± 0.3 | -098.70 |  |
| V3 | 42.3 ± 2.0 | 24.4 ± 0.3 | -190.22 | + |
| V4 | 41.6 ± 1.2 | 24.4 ± 0.2 | -221.10 | ++ |
| V5 | 47.8 ± 1.7 | 25.2 ± 0.4 | -072.33 |  |
| V6 | 43.7 ± 1.5 | 24.8 ± 0.3 | -137.22 | + |
| V7 | 41.2 ± 2.1 | 24.8 ± 0.3 | -218.22 | ++ |
| V8 | 44.8 ± 2.0 | 24.8 ± 0.2 | -121.34 |  |
| V9 | 42.4 ± 1.6 | 24.4 ± 0.3 | -107.63 | + |
| V10 | 47.9 ± 2.7 | 24.9 ± 0.3 | -087.71 |  |
| **Food Library** |  |  |  |  |
| F1 | 49.6 ± 1.4 | 25.6 ± 0.2 | -33.25 |  |
| F2 | 49.1 ± 1.7 | 25.5 ± 0.2 | -37.22 |  |
| F3 | 48.7 ± 1.9 | 25.5 ± 0.2 | -54.85 |  |
| F4 | 49.0 ± 1.3 | 25.3 ± 0.2 | -40.71 |  |
| F5 | 47.5 ± 1.4 | 25.5 ± 0.2 | -77.20 |  |
| F6 | 47.7 ± 1.5 | 25.0 ± 0.2 | -72.53 |  |
| F7 | 46.5 ± 2.0 | 25.3 ± 0.3 | -42.12 |  |
| F8 | 45.5 ± 2.8 | 24.4 ± 0.2 | -93.34 | ? |
| F9 | 48.6 ± 1.8 | 25.2 ± 0.2 | -39.16 |  |
| F10 | 46.8 ± 1.7 | 25.7 ± 0.2 | -29.80 |  |
| *Simulations done at pH 5.7* |  |  |  |  |
| Receptor [Av. 5MN2] |  |  |  |  |
| D4_low | 41.2 ± 1.7 | 24.3 ± 0.2 | -176.03 | ++ |
| V4_low | 43.5 ± 1.6 | 24.6 ± 0.2 | -156.85 | ++ |
| V7_low | 43.7 ± 1.6 | 24.7 ± 0.2 | -148.15 | ++ |

The variation in the values of the molecular descriptors, θ and δ, characterizing the interdomain gap between the ectodomains of FcγRIIIa as complex to the selected ligands are plotted in **Figure 3**. The left panels show the values from simulations carried out at theoretical pH of 7.4. For all three complexes, the average values of θ and dis were lower than the values observed for the FcγRIIIa receptor alone (please see the dotted black line as reference). Additionally, the calculated R_g_ values for the complexes were much lower than the R_g_ values computed for the abstracted receptor. This indicated that the three small ligands can significantly close the gap between the two ectodomains of FcγRIIIa in the simulation timelines. We also performed calculations of these complexes at pH 5.7 to see any influence on the conformational properties and/or any unexpected opening of the receptor domains at pH conditions representing the endosome. The right panels in **Figure 3** show the variance in the values of molecular descriptors of FcγRIIIa and R_g_ of the complexes at pH 5.7. The interdomain angle between the two ectodomains of receptor was slightly more open at lower pH, but values were within the variation seen during the simulation. The other parameters, δ and R_g_ remained similar like those computed for structures simulated at pH 7.4. These results indicate that the three lead ligands, D4, V4 and V7 will not allow the ectodomains of FcγRIIIa to open in the low pH environment of the endosomes and may remain bound to the receptor through the entry and exit cycle(s). **Figure 4** shows the snapshots of the ligands in their average pose on the FcγRIIIa receptor at pH 7.4 and 5.7. For each ligand, the pose appears to be similar at both the pH values. Since this work was carried out to explore a hypothesis, over-analysis of interactions was skipped in absence of experimental biophysical evidence that identified lead compounds can hold the two ectodomains of FcγRIIIa receptor tightly closed.

**Figure 3.**
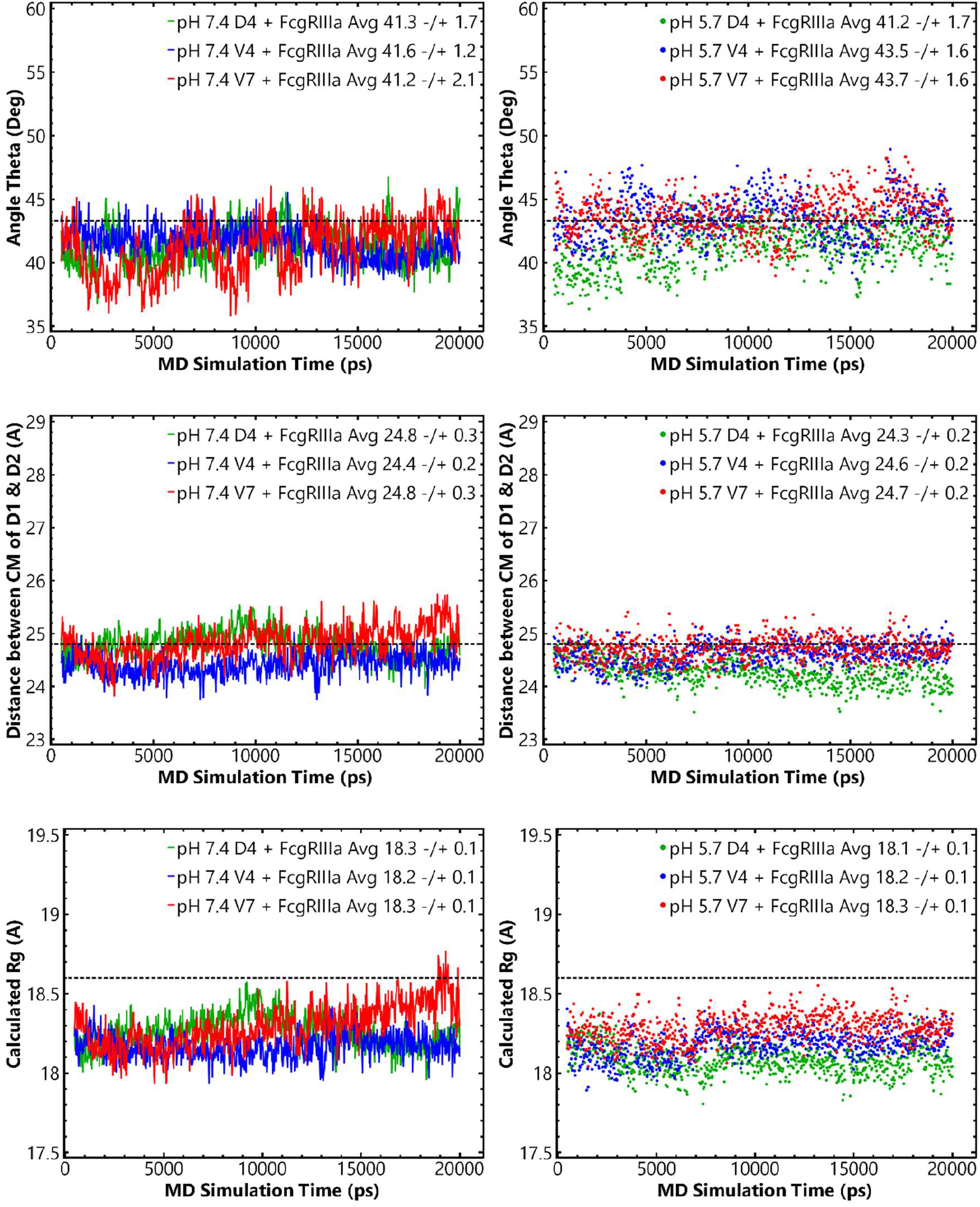
Variation in the calculated values of θ and δ in the simulated structures of the complexes of selected ligands with FcγRIIIa as a function of the MD simulation time at pH 7.4 and 5.7 are presented here. In the legends of the plots, the starting structure of the calculation, average and standard deviation of the calculated descriptor are mentioned. The dotted horizontal line in each panel represents the average value for the unliganded receptor from its MD simulations.

**Figure 4.**
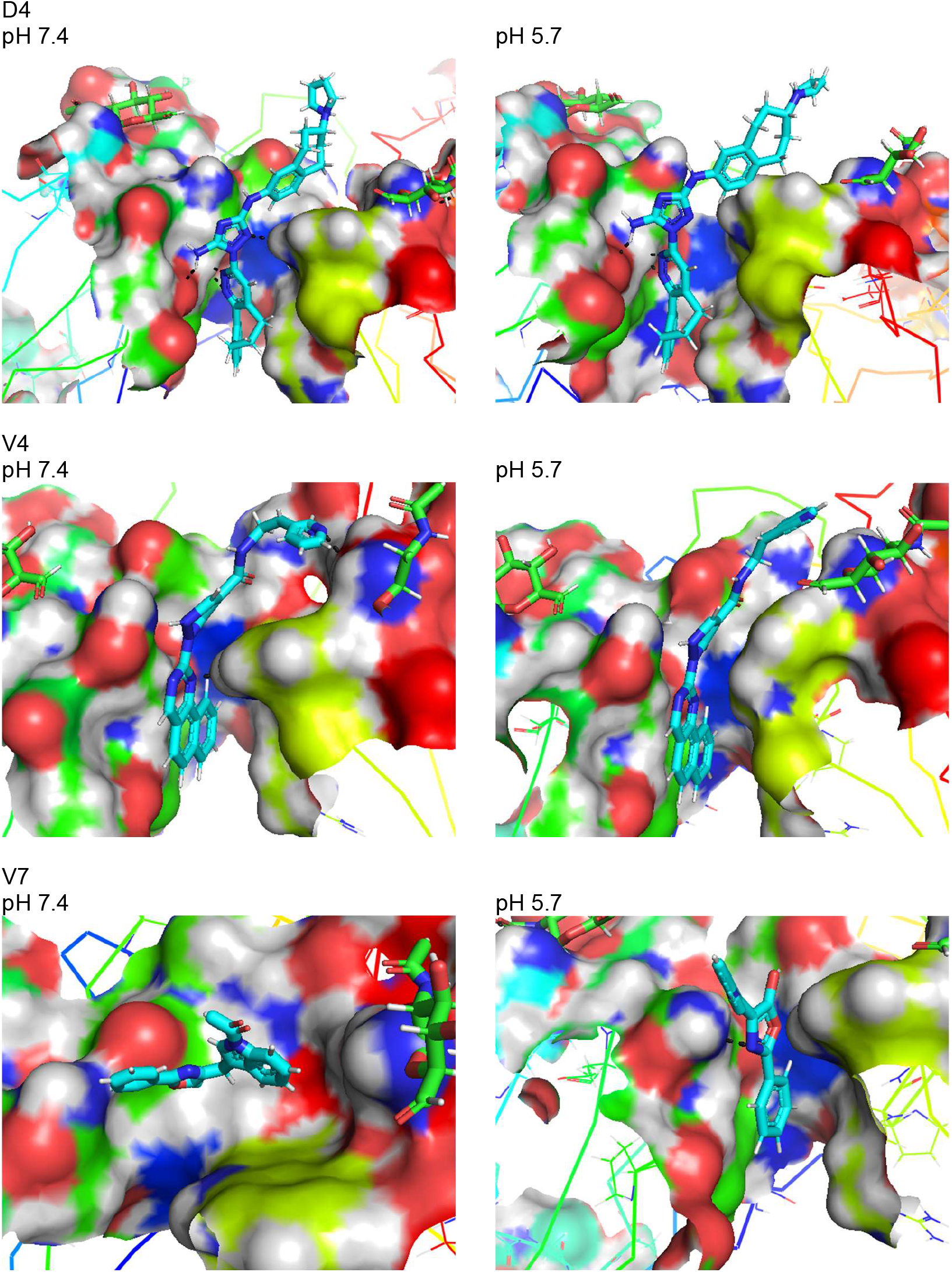
The average pose of the drug-like molecules D4, V4 and V7 between the cleft of the two ectodomains of FcγRIIIa during the MD simulations of their respective complexes at pH 7.4 and 5.7 are shown here. The drugs are presented as cyan, blue and red sticks with interacting surface of the FcγRIIIa receptor are shown as solid surface.

### FcγRIIIb and its reactivity to selected small ligands

Query remained if the selected ligands would cross-react with the ectodomains of the other receptor i.e. FcγRIIIb and induce structural changes as it does for FcγRIIIa, as per our simulations? For this purpose, first we simulated three structures of FcγRIIIb as described in the method section. Starting with the PDB ID 6EAQ [20], three structures of ectodomains of FcγRIIIb were extracted for MD simulations: aglycosylated or protein only structure, glycosylated structure and glycosylated structure bound to Fc portion as in the crystal structure. MD simulations were done only at pH 7.4, because that is where the ligands might interact first with this receptor. The molecular descriptors, θ and δ were calculated for the FcγRIIIb and are plotted as a function of simulation time in **Figure 5**. Interdomain angles and distance between the center of masses were higher for the glycosylated receptor without or with Fc portion. The protein only structure showed decrement in the descriptors values like that observed for FcγRIIIa indicating that the sugar moieties in crystal structure and Fc influence the ectodomain architecture of the FcγRIIIb receptor. Compared to FcγRIIIa, FcγRIIIb reflected more open structure of the acute angle formed by the ectodomains. The calculated R_g_ values for the protein only and glycosylated structure of FcγRIIIb indicated no significant difference which implied that for our study, the molecular descriptors are better track of conformational changes of interest (**Supplementary Figure S5**). Next, MD simulations of the complexes of glycosylated FcγRIIIb with the selected ligands were done at pH 7.4 and the molecular descriptors were tracked (**Right Panels in Figure 5**). The average values of θ, δ and binding energies are mentioned in the table below the panels. As can be seen, D4 could not keep the two ectodomains closer than the calculated for the receptor alone. While the interdomain angle was relatively lower for the complexes with V4 and V7, δ values were higher than that observed for the FcγRIIIb receptor alone. The computed binding energies of these ligands to FcγRIIIb receptor were substantially lower when compared to the values calculated for FcγRIIIa at both pH values. Overall analysis reflected that these ligands may bind very weakly or not at all to FcγRIIIb receptor. Of course, wet lab experiments will be the reliable answer on this aspect.

**Figure 5.**
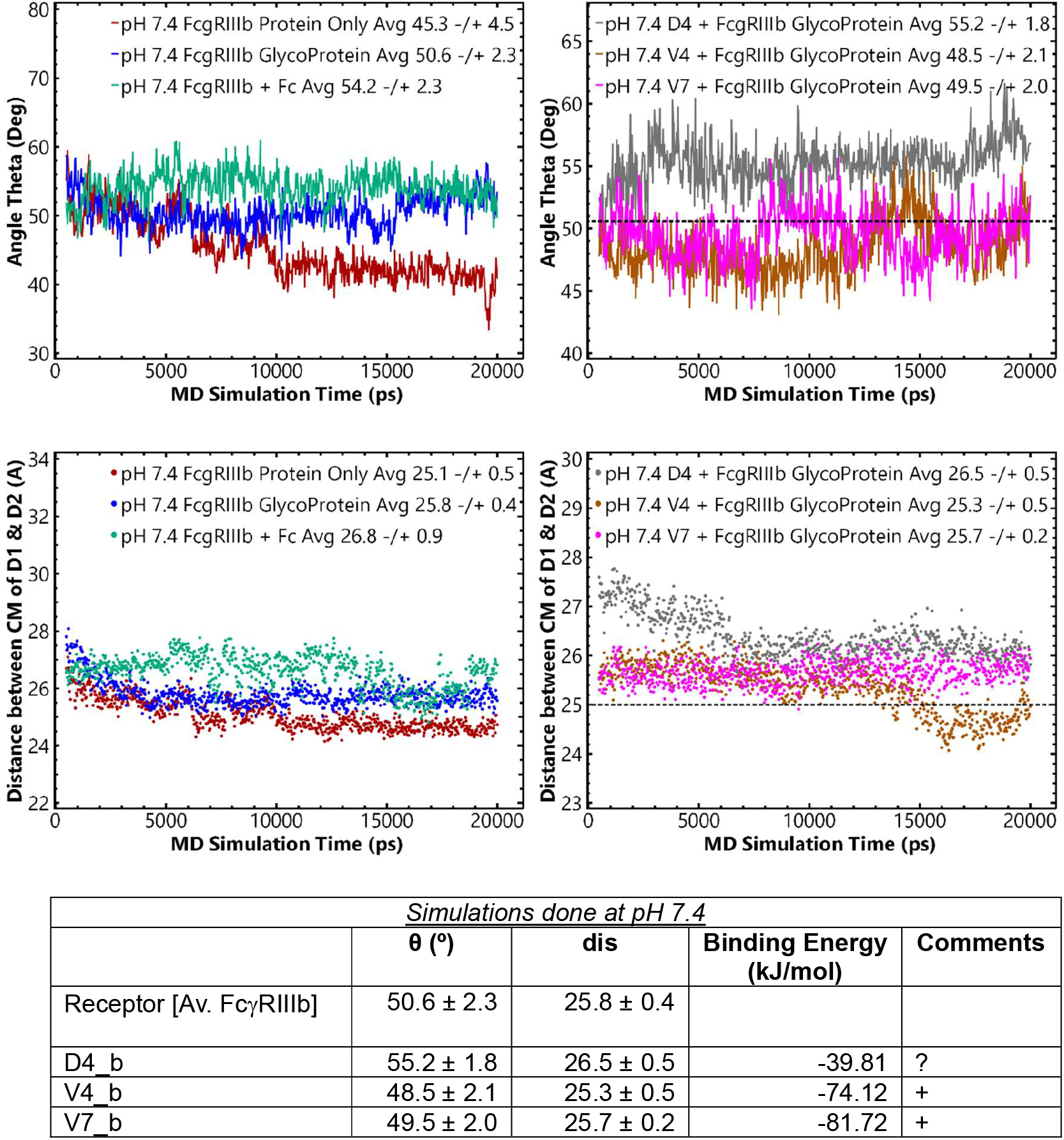
Variation in the calculated values of θ and δ in the simulated structures of the FcγRIIIb -/+ binding partners are plotted as a function of the MD simulation time at pH 7.4 are presented here. In the legends of the plots, the starting structure of the calculation, average and standard deviation of the calculated descriptor are mentioned. The dotted horizontal line in the right panels represents the average value for the unliganded receptor from its MD simulations. The table below the panels sums up the descriptor and binding energy values with the three selected ligands.

## Conclusions

This computational work was initiated to explore a hypothesis if small molecules can dock between the ectodomains of FcγRIIIa and keep them closed like the two reported affimers which may be therapeutic against ITP [6]. Our docking and modeling experiments led to the identification of three molecules which we named D4, V4 and V7. Tracking the molecular descriptor values suggest that these three molecules would bind to FcγRIIIa and will tightly close its two ectodomains. In their binding induced closed state, FcγRIIIa will be rendered allosterically incapable of binding subsequently to Fc portion of IgG antibodies. The smiles code for the three selected molecules are: C1CCN(C1)[C@H]2CCC3=C(CC2)C=C(C=C3)NC4=NN(C(=N4)N)C5=NN=C6C(=C5)CCCC7=CC=C C=C76; Cc1c(cnn1c1ncc2CCc3ccccc3c2n1)C(=O)NCCc1cccnc1 and C(c1cn(C(=O)C)c2ccccc12)c1c(O)oc(c2ccccc2)n1. Searches provided that D4 is also known as Bemcentinib (R428) and is a potent and selective inhibitor of Axl with an IC50 of 14 nM [23] and some other relevant indicators [24,25], but none directly or indirectly correlating with ITP. Unfortunately, no information could be found for any bio-functionality for V4 and V7. Before proceeding further with these ligands or their analogs, experimental validation about their ability to keep the FcγRIIIa receptor closed needs to be verified. For requisite experiments, the key reagents like purified ectodomains of FcγRIIIa with native-like glycosylation, its mutants and ligands appear achievable. Biophysical experiments like small angle X-ray scattering (SAXS) can easily and reliably provide tracking of conformational properties of FcγRIIIa as a function of buffer pH, temperature and ratio of ligand available for binding [26,27]. Experimental binding energies can be evaluated by different techniques like SPR, Isothermal calorimetry (ITC) or Biolayer interference (BLI) etc. Confirming the ability of these ligands (and their analogs) to bind tightly FcγRIIIa, and induce its ectodomains to remain closed will support their case to be further evaluated by cell-line based experiments as mentioned before [6]. Availability of reliable animal models of human ITP including humanized mice would aid in furthering translation potential of this line of thinking. In the Supplementary material, three GROK3 (www.grok.com) based search results have been summarized which detail the incidence to mortality rates in different countries from ITP, drugs under clinical evaluations and different clinical trials underway (**Supplementary Information 1**). (*Readers can also perform their own surveys and summarizations*). There are some small molecule inhibitors of downstream kinases which are under clinical trials for chronic ITP and are showing promising results in increasing the plated counts in sustained manner (**Supplementary Information 2**). **Supplementary Information 3** briefly describes the molecules under clinical trials underway for ITP. It Is pertinent to mention here that for pediatric ITP cases, the first line of non-steroidal therapies against ITP are transfusion of platelets and/or injectable IvIG. Both are usually fractioned from human plasma and undergo rigorous testing and processing before being given to the patients. Even the reported affimers, being proteins will have to undergo the challenges of overproduction to purification to storage conditions, before they can be given as injectable therapies or some other innovative way. Small molecule interventions do hold better translational potential due to lower cost of productions and range of formulations are available for oral deliveries. As mentioned, few small molecule inhibitors of downstream kinases are in clinical trials, molecules working on our hypothesis would be more specific and may show less off-target effects especially under chronic conditions requiring prolonged therapies. Molecules blocking this pathway could be viable co-therapies too. Our lab will continue to dig deeper into this hypothesis, and other interested researchers are welcome to explore with us.

## Acknowledgements

Author acknowledges availability of open-source databases from NCBI and RCSB for unfettered access. This work was conceptualized and executed while the author was on extended leave.

## References

1. Hamdan TA, Lang PA, Lang KS. The Diverse Functions of the Ubiquitous Fcgamma Receptors and Their Unique Constituent, FcRgamma Subunit. Pathogens. 2020 Feb 20;9(2).

2. Junker F, Gordon J, Qureshi O. Fc Gamma Receptors and Their Role in Antigen Uptake, Presentation, and T Cell Activation. Front Immunol. 2020;11:1393.

3. Nimmerjahn F, Ravetch JV. Fcgamma receptors as regulators of immune responses. Nat Rev Immunol. 2008 Jan;8(1):34–47.

4. Qiu WQ, de Bruin D, Brownstein BH, et al. Organization of the human and mouse low-affinity Fc gamma R genes: duplication and recombination. Science. 1990 May 11;248(4956):732–5.

5. Machado LR, Hardwick RJ, Bowdrey J, et al. Evolutionary history of copy-number-variable locus for the low-affinity Fcgamma receptor: mutation rate, autoimmune disease, and the legacy of helminth infection. Am J Hum Genet. 2012 Jun 8;90(6):973–85.

6. Robinson JI, Baxter EW, Owen RL, et al. Affimer proteins inhibit immune complex binding to FcgammaRIIIa with high specificity through competitive and allosteric modes of action. Proc Natl Acad Sci U S A. 2018 Jan 2;115(1):E72–E81.

7. Clarkson SB, Bussel JB, Kimberly RP, et al. Treatment of refractory immune thrombocytopenic purpura with an anti-Fc gamma-receptor antibody. N Engl J Med. 1986 May 8;314(19):1236–9.

8. Bosques CJ, Manning AM. Fc-gamma receptors: Attractive targets for autoimmune drug discovery searching for intelligent therapeutic designs. Autoimmun Rev. 2016 Nov;15(11):1081–1088.

9. MacFarlane LA, Todd DJ. Kinase inhibitors: the next generation of therapies in the treatment of rheumatoid arthritis. Int J Rheum Dis. 2014 May;17(4):359–68.

10. Blazquez-Moreno A, Park S, Im W, et al. Transmembrane features governing Fc receptor CD16A assembly with CD16A signaling adaptor molecules. Proc Natl Acad Sci U S A. 2017 Jul 11;114(28):E5645–E5654.

11. DiLillo DJ, Tan GS, Palese P, et al. Broadly neutralizing hemagglutinin stalk-specific antibodies require FcgammaR interactions for protection against influenza virus in vivo. Nat Med. 2014 Feb;20(2):143–51.

12. Jing Y, Ni Z, Wu J, et al. Identification of an ADAM17 cleavage region in human CD16 (FcgammaRIII) and the engineering of a non-cleavable version of the receptor in NK cells. PLoS One. 2015;10(3):e0121788.

13. Lee J, Zhang T, Hwang I, et al. Epigenetic modification and antibody-dependent expansion of memory-like NK cells in human cytomegalovirus-infected individuals. Immunity. 2015 Mar 17;42(3):431–42.

14. Wang TT, Sewatanon J, Memoli MJ, et al. IgG antibodies to dengue enhanced for FcgammaRIIIA binding determine disease severity. Science. 2017 Jan 27;355(6323):395–398.

15. Land H, Humble MS. YASARA: A Tool to Obtain Structural Guidance in Biocatalytic Investigations. Methods Mol Biol. 2018;1685:43–67.

16. Ozvoldik K, Stockner T, Krieger E. YASARA Model-Interactive Molecular Modeling from Two Dimensions to Virtual Realities. J Chem Inf Model. 2023 Oct 23;63(20):6177–6182.

17. Dey M, Gupta A, Badmalia MD, et al. Visualizing gaussian-chain like structural models of human alpha-synuclein in monomeric pre-fibrillar state: Solution SAXS data and modeling analysis. Int J Biol Macromol. 2025 Feb;288:138614.

18. Kalidas N, Peddada N, Pandey K, et al. SAXS data based glycosylated models of human alpha-1-acid glycorprotein, a key player in health, disease and drug circulation. J Biomol Struct Dyn. 2025 Mar 8:1–15.

19. Labbe CM, Rey J, Lagorce D, et al. MTiOpenScreen: a web server for structure-based virtual screening. Nucleic Acids Res. 2015 Jul 1;43(W1):W448–54.

20. Roberts JT, Barb AW. A single amino acid distorts the Fc gamma receptor IIIb/CD16b structure upon binding immunoglobulin G1 and reduces affinity relative to CD16a. J Biol Chem. 2018 Dec 21;293(51):19899–19908.

21. Waterhouse A, Bertoni M, Bienert S, et al. SWISS-MODEL: homology modelling of protein structures and complexes. Nucleic Acids Res. 2018 Jul 2;46(W1):W296-W303.

22. Studer G, Rempfer C, Waterhouse AM, et al. QMEANDisCo-distance constraints applied on model quality estimation. Bioinformatics. 2020 Mar 1;36(6):1765–1771.

23. Hoel A, Osman T, Hoel F, et al. Axl-inhibitor bemcentinib alleviates mitochondrial dysfunction in the unilateral ureter obstruction murine model. J Cell Mol Med. 2021 Aug;25(15):7407–7417.

24. Loges S, Heuser M, Chromik J, et al. Bemcentinib as monotherapy and in combination with low-dose cytarabine in acute myeloid leukemia patients unfit for intensive chemotherapy: a phase 1b/2a trial. Nat Commun. 2025 Mar 23;16(1):2846.

25. Wu KM, Xu QH, Liu YQ, et al. Neuronal FAM171A2 mediates alpha-synuclein fibril uptake and drives Parkinson’s disease. Science. 2025 Feb 21;387(6736):892–900.

26. Sagar A, Haleem N, Bashir YM, et al. Search for non-lactam inhibitors of mtb beta-lactamase led to its open shape in apo state: new concept for antibiotic design. Sci Rep. 2017 Jul 24;7(1):6204.

27. Sagar A, Arif E, Solanki AK, et al. Targeting Neph1 and ZO-1 protein-protein interaction in podocytes prevents podocyte injury and preserves glomerular filtration function. Sci Rep. 2017 Sep 21;7(1):12047.

