## Supplementary Material for "SMALL MOLECULES WHICH CAN KEEP THE ECTODOMAINS OF HUMAN FcγRIIIa “CLOSED”: THERAPY FOR AUTOIMMUNE THROMBOCYTOPENIA PURPURA?"

### Supplementary Figure S1

CLUSTAL O (1.2.4) multiple sequence alignment*

```
sp|P08637|FCGIIIA_HUMAN      MWQLLLPTALLLLVSAGMRTEDLPKAVVFLEPQWYRVLEKDSVTLKCQGAYSPEDNSTQW 60
sp|O75015|FCGIIIB_HUMAN      MWQLLLPTALLLLVSAGMRTEDLPKAVVFLEPQWYSVLEKDSVTLKCQGAYSPEDNSTQW 60
                             *********************************** ************************

sp|P08637|FCGIIIA_HUMAN      FHNESLISSQASSYFIDAATVDDSGEYRCQTNLSTLSDPVQLEVHIGWLLLQAPRWVFKE 120
sp|O75015|FCGIIIB_HUMAN      FHNENLISSQASSYFIDAATVNDSGEYRCQTNLSTLSDPVQLEVHIGWLLLQAPRWVFKE 120
                             ****.****************:**************************************

sp|P08637|FCGIIIA_HUMAN      EDPIHLRCHSWKNTALHKVTYLQNGKGRKYFHHNSDFYIPKATLKDSGSYFCRGLFGSKN 180
sp|O75015|FCGIIIB_HUMAN      EDPIHLRCHSWKNTALHKVTYLQNGKDRKYFHHNSDFHIPKATLKDSGSYFCRGLVGSKN 180
                             **************************.**********:*****************.****

sp|P08637|FCGIIIA_HUMAN      VSSETVNITITQGLAVSTISSFFPPGYQVSFCLVMVLLFAVDTGLYFSVKTNIRSSTRDW 240
sp|O75015|FCGIIIB_HUMAN      VSSETVNITITQGLAVSTISSFSPPGYQVSFCLVMVLLFAVDTGLYFSVKTNI------- 233
                             ********************** ******************************

sp|P08637|FCGIIIA_HUMAN      KDHKFKWRKDPQDK  254
sp|O75015|FCGIIIB_HUMAN      --------------  233
```

*Ectodomains have 207 residues Including the signal sequence at N-terminal; Yellow highlighted residues were predicted to form transmembrane α-helix; Gly129 shown to be critical in high affinity binding to Fc of IgG1 is shown with an arrow above it (15).

Alignment of the primary structures of human FcγRIIIa and FcγRIIIb Receptors are shown above. UniProt accession numbers are mentioned on the right.

**Supplementary Figure S2**

A	Schematic representation of how the antigen bound IgG1 will interact with the ectodomains of FcγRIIIa via latter's Fc portion outside the cells is presented below. The whole complex is internalized via pinocytosis, and if released from the FcγRIIIa receptor, then the released IgG and antigen undergoes degradation at lower pH in the endosomes. Since bound antibodies and their antigens are protected from degradation this could be the mechanism by which high affinity auto antibodies and antigens can survive and come out again.

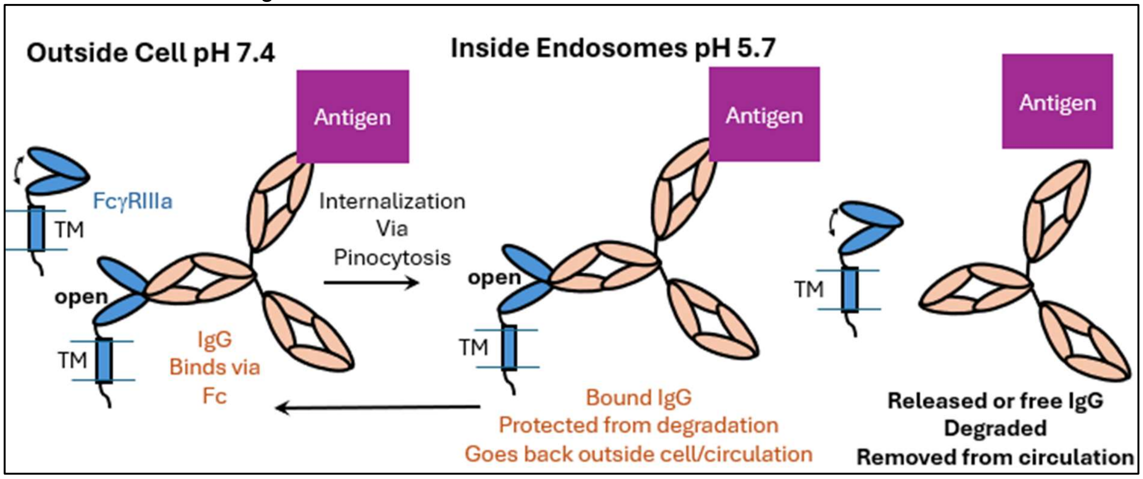


B	Schematic representation of how IvIG molecules injected during therapies bind to ectodomains of FcγRIIIa and competitivity block the binding of antigen bound antibodies. As complex, IvIG are internalized, not degraded and they may come out from the endosomes. On the other hand, antibodies and antigens as free or complex enter the endosomes by pinocytosis unassisted by FcγRIIIa, they get degraded and are taken out of the circulation, including auto antibodies.

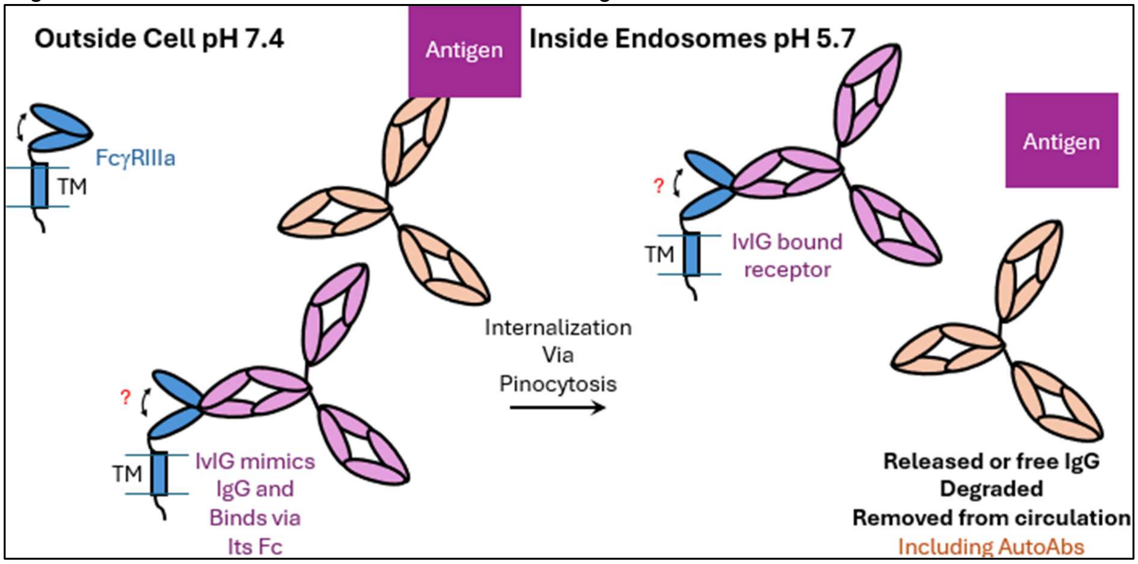

C Schematic representation of how isolated Fc portion of IgG1 or its bio-mimics will bind to ectodomains of FcγRIIIa and block the binding of antigen bound antibodies. As complex, these molecules get internalized, not degraded and if not released by FcγRIIIa receptor, they may come out from the endosomes. On the other hand, antibodies and antigens as free or complex enter the endosomes by pinocytosis unassisted by FcγRIIIa, they get degraded and are taken out of the circulation, including auto antibodies.

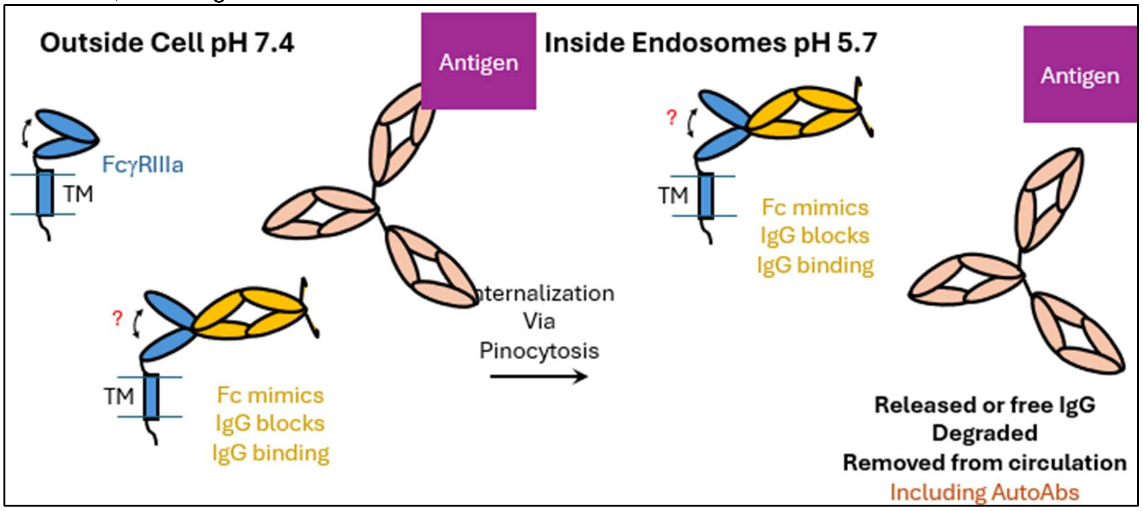


D Schematic representation of how designer affimers AfF4 and AfG3 will bind to ectodomains of FcγRIIIa and block the binding of antigen bound antibodies. As described, AfF4 interferes with binding of Fc portion and AfG3 achieves allosteric inhibition. Very likely, as complexed to FcγRIIIa, these affimers will get internalized, not degraded and may come out from the endosomes. On the other hand, antibodies and antigens as free or complex enter the endosomes by pinocytosis unassisted by FcγRIIIa, they get degraded and are taken out of the circulation, including auto antibodies.

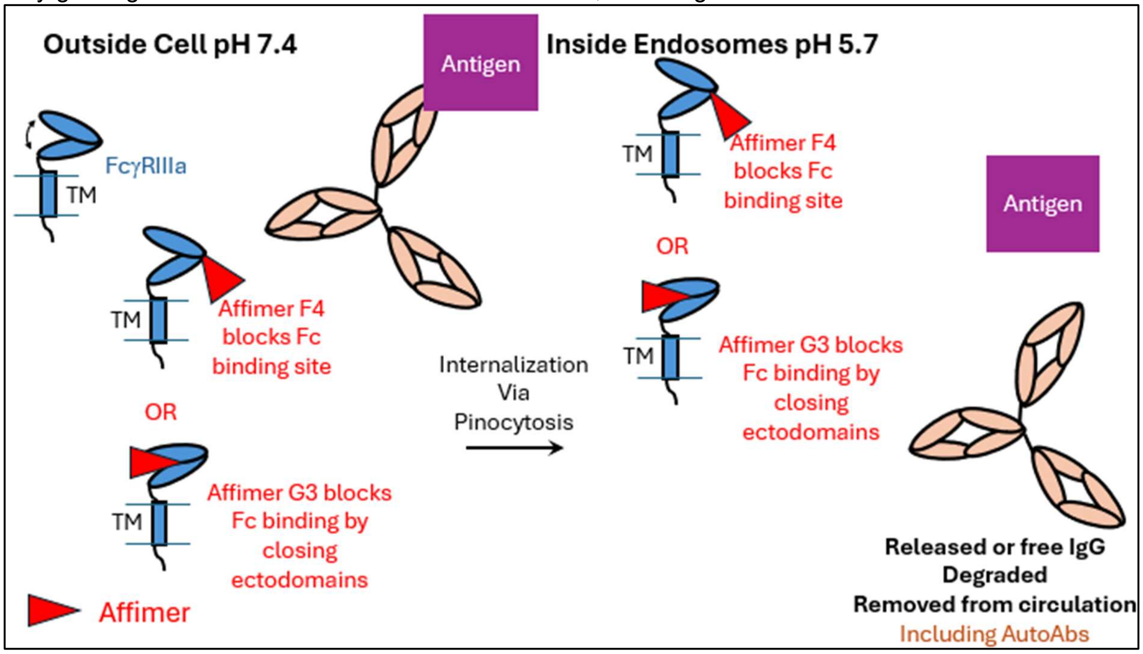

E Schematic representation of how our planned small molecules will bind to ectodomains of FcγRIIIa and allosterically block the binding of antigen bound antibodies. We postulate that complexed to FcγRIIIa, our molecules will get internalized, not get degraded and may come out from the endosomes bound to FcγRIIIa receptor. On the other hand, antibodies and antigens as free or complex enter the endosomes by pinocytosis unassisted by FcγRIIIa, they get degraded and are taken out of the circulation, including auto antibodies.

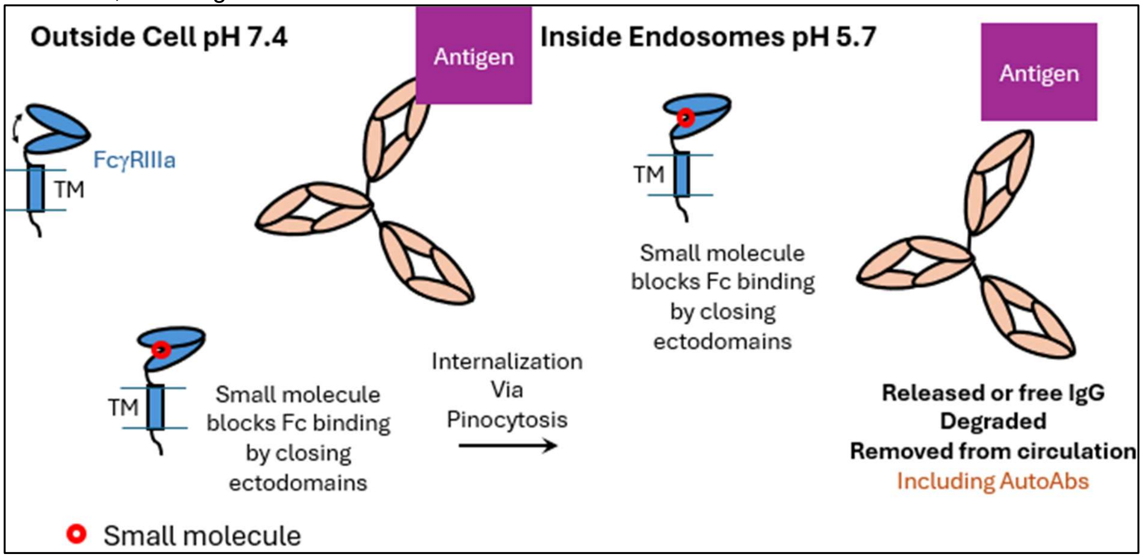

**Supplementary Figure S3**

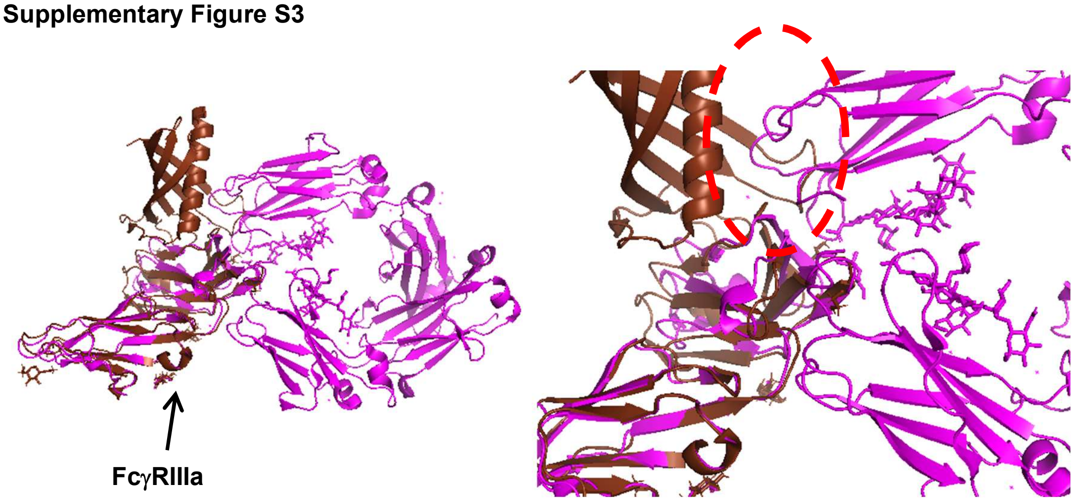


The coordinates of the FcγRIIIa chain in the crystal structures of complexes of FcγRIIIa with affimer AfF4 and Fc portion of IgG1 are superimposed here (PDB IDs 5MNL9 brown cartoon and 7URU magenta cartoon, respectively). Aligned FcγRIIIa chains in the two crystal structures are pointed by an arrow in the left panel. The right image shows a zoom-in of the interaction area, and the conflicting residues or loops in the structures of the bound affimer and Fc are shown with the red dashed circle.

**Supplementary Figure S4**

Results from MD simulations at pH 7.4

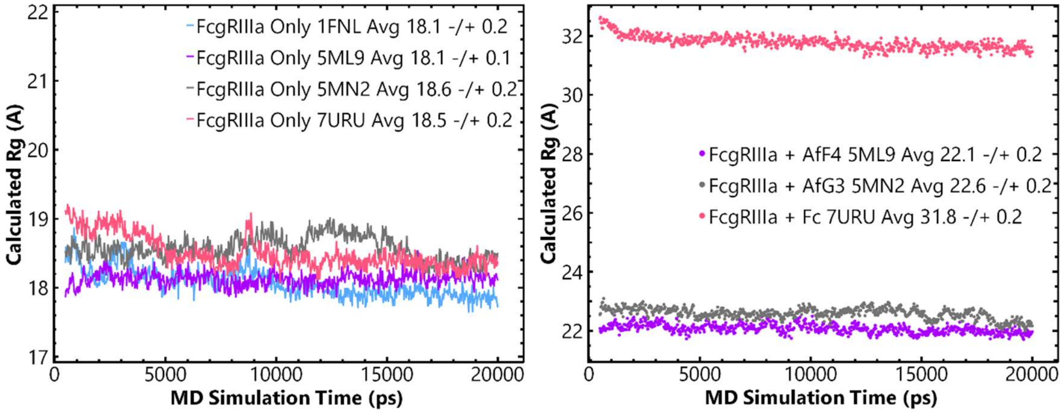


Results from MD simulations at pH 5.7

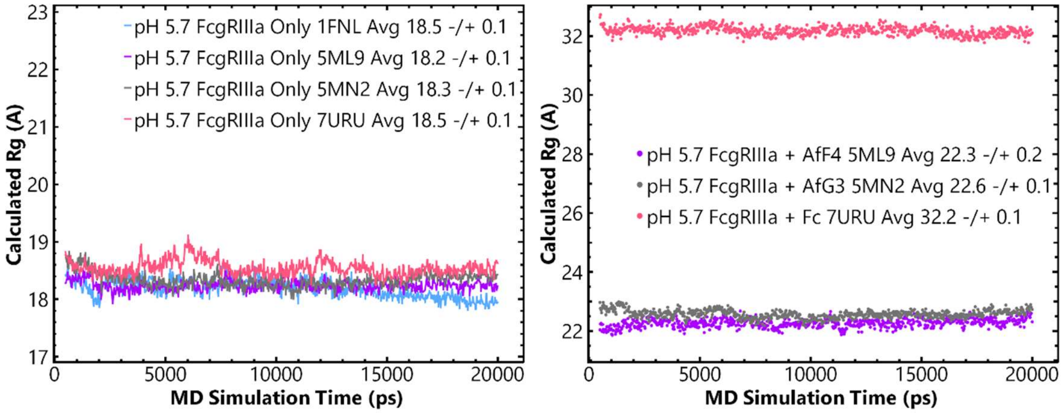


**Variation in the calculated Rg values of the simulated structures of FcγRIIIa -/+ binding partners as a function of the MD simulation time at pH 7.4 and 5.7 are presented here.** In the legends of the plots, the starting structure of the simulation, average and standard deviation of the calculated descriptor are mentioned.

**Supplementary Table ST1**

The short-listed molecules from docking of **Approved Drug library**

| Given Name | Name ZINC Database ID |
|---|---|
| D1 | Metergotamine_ZINC000072266819 |
| D2 | R428_ZINC000051951669 |
| D3 | Ergotamine_ZINC000052955754 |
| D4 | R428_ZINC000051951668 |
| D5 | Dihydroergotamine_ZINC000003978005 |
| D6 | Disogluside_ZINC000008214547 |
| D7 | Venetoclax_ZINC000150338755 |
| D8 | Losulazine_ZINC000004216779 |
| D9 | Mk3207_ZINC000103760984 |
| D10 | Bolazine_ZINC000008214506 |

The short-listed molecules from docking of **Diverse library**

| Given Name | Smiles | Library ID [MTiOpenScreen] |
|---|---|---|
| V1 | [CH2-2]1[CH2-2][CH2-2]N([CH2-2]1)C(=O)N1[C-]2[CH2-2][CH2-2][C-]1C(=C([CH2-2]2)c1[c-][c-]c(c([c-]1)F)c1[c-][c-][c-][c-][c-]1)C(=O)O[CH3-3] | 85148520_Intermediate |
| V2 | N(S(=O)(=O)c1[c-]c2[c-][c-]n(c2[c-][c-]1)C(=O)[CH3-3])[CH2-2][CH2-2]C(=O)N[C-]1[CH2-2][CH2-2][CH2-2]c2[c-][c-][c-][c-]c12 | 26724445_Intermediate |
| V3 | [CH2-2]1[C-](Oc2c1[c-]c([c-]c2F)c1[c-]c2[c-][c-][c-][c-]c2n[c-]1)[CH2-2]NC(=O)c1[c-]nsn1 | 124948075_Intermediate |
| V4 | [CH3-3]c1c([c-]nn1c1n[c-]c2[CH2-2][CH2-2]c3[c-][c-][c-][c-]c3c2n1)C(=O)N[CH2-2][CH2-2]c1[c-][c-][c-]n[c-]1 | 124948465_Accepted |
| V5 | c1([c-][c-][c-]c(C#N)[c-]1)NC(=O)[CH2-2]Sc1nc(=O)c2c([c-]sc2[nH]1)c1[c-][c-][c-][c-][c-]1 | 85176827_Intermediate |
| V6 | [CH2-2]1[CH2-2]N([CH2-2][CH2-2]N1C(=O)c1[c-][c-]c([c-][c-]1)c1[c-][c-][c-][c-][c-]1)C(=O)c1[c-][c-][c-]s1 | 17464346_Intermediate |
| V7 | [CH2-2](c1[c-]n(C(=O)[CH3-3])c2[c-][c-][c-][c-]c12)c1c([O-])oc(c2[c-][c-][c-][c-][c-]2)n1 | 17431512_Intermediate |
| V8 | c1([c-][c-]c([c-][c-]1)NC(=O)c1[c-][c-][c-]o1)C(=O)N[CH2-2][C-]1[CH2-2]Oc2[c-][c-][c-][c-]c2O1 | 4263243_Intermediate |
| V9 | [C-]1(C(=C(Oc2c1c(=O)[nH]c1[c-][c-][c-][c-]c21)N)C#N)c1[c-][c-][c-][c-]c1C(F)(F)F | 7996409_Intermediate |
| V10 | C(=O)(N1[CH2-2][CH2-2]c2nnc(n2[CH2-2][CH2-2]1)[CH2-2]NC(=O)c1[c-][c-][c-]c([c-]1)F)[CH2-2][CH2-2]c1[c-][c-][c-][c-][c-]1 | 124947978_Intermediate |

The short-listed molecules from docking of **Food library**

| Given Name | Name ZINC Database ID |
|---|---|
| F1 | beta_Amyrenone_ZINC000031165761 |
| F2 | Oleanolic_acid_ZINC000003881983 |
| F3 | 3_Oxo_olean_12_en_28_oic_acid_ZINC000013558220 |
| F4 | 2alpha_3alpha_Dihydroxyolean_12_en_28_oic_acid_ZINC000008952016 |
| F5 | Taraxasterol_acetate_ZINC000169748443 |
| F6 | 9_11__Dehydroglycyrrhetic_acid_ZINC000012871005 |
| F7 | Katonic_acid_ZINC000014980278 |
| F8 | Sapogenins_ZINC000257348756 |
| F9 | Ursolic_acid_ZINC000004273371 |
| F10 | Soyasapogenol_B_1_ZINC000247952264 |

**Supplementary Figure S5**

Results from MD simulations at pH 7.4

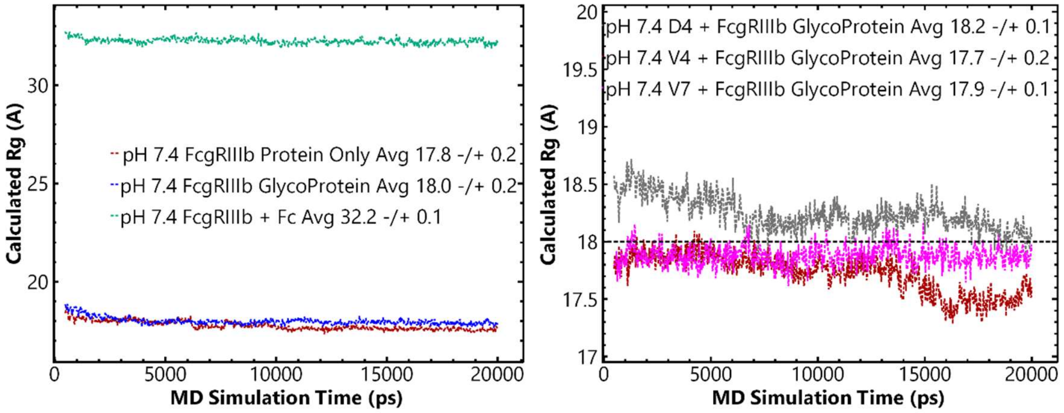


**Variation in the calculated Rg values of the simulated structures of FcγRIIIb -/+ binding partners as a function of the MD simulation time at pH 7.4 are presented here.** In the legends of the plots, the starting structure of the simulation, average and standard deviation of the calculated descriptor are mentioned.

**Supplementary Information 1**
**GROK3 based research summary on molecules in clinical trials for ITP**
*Done on March 21, 2025 7:58 AM CST*

This table includes new countries across different continents to broaden the scope, with population estimates for 2025 and ITP metrics based on studies or reasonable extrapolations. Estimates are marked where data is inferred.

#### Further Expanded Country-Wise Data on ITP with Age Breakdown

| **Country** | **Population (2025)** | **Age Group** | **Incidence Rate (per 100,000)** | **Prevalence (per 100,000)** | **Estimated Cases (Incidence)** | **Mortality Rate (per 100,000)** | **Common Treatments** | **Notes** |
|---|---|---|---|---|---|---|---|---|
| **United States** | 345,000,000 | Children | 4-5 | 5-10 | 2,760-3,450 | 0.1-0.2 | Observation, IVIG | Acute, often post-viral |
| | | Adults | 2-3 | 12-25 | 5,520-8,280 | 0.2-0.5 | Steroids, TPO-RA, Splenectomy | Chronic in 70-80% of cases |
| **United Kingdom** | 68,000,000 | Children | 3-4 | 5-8 | 408-544 | 0.1 | Observation, Steroids | High spontaneous remission |
| | | Adults | 2-3 | 15-25 | 1,088-1,632 | 0.2-0.3 | Rituximab, Splenectomy | UK GPRD data |
| **Japan** | 123,000,000 | Children | 3-5 | 5-10 | 738-1,230 | 0.1 | Observation, H. pylori Rx | H. pylori link in some cases |
| | | Adults | 2-4 | 15-25 | 2,460-4,920 | 0.1-0.2 | Steroids, Rituximab | Aging population impact |
| **India** | 1,428,000,000 | Children | 2-4 (est.) | 5-10 (est.) | 5,712-11,424 | Unknown | Observation, Steroids | Limited pediatric studies |
| | | Adults | 1-3 (est.) | 5-15 (est.) | 14,280-42,840 | Unknown | Steroids, Observation | Underreported chronic cases |
| **Brazil** | 203,000,000 | Children | 3-5 (est.) | 5-10 (est.) | 1,218-2,030 | Unknown | IVIG, Observation | Acute cases predominate |
| | | Adults | 2-4 (est.) | 10-15 (est.) | 2,436-4,872 | Unknown | Steroids, IVIG | Emerging adult data |
| **Canada** | 41,000,000 | Children | 4-6 (est.) | 5-10 (est.) | 328-492 | 0.1-0.2 | Observation, IVIG | Similar to US patterns |
| | | Adults | 2-4 | 10-20 | 656-1,312 | 0.2-0.4 | Steroids, TPO-RA | North American trends |
| **Germany** | 84,000,000 | Children | 3-5 | 5-10 | 504-840 | 0.1 | Observation, Steroids | Nordic/EU registry data |
| | | Adults | 2-4 | 10-18 | 1,680- | | | |

| | | | | | | | | |
|---|---|---|---|---|---|---|---|---|
| | | | | | 3,360 | 0.1-0.3 | TPO-RA, Splenectomy | Chronicity common in adults |
| **China** | 1,425,000,000 | Children | 2-4 (est.) | 5-10 (est.) | 5,700-11,400 | Unknown | Observation, Steroids | Sparse data, large population |
| | | Adults | 1-3 (est.) | 5-15 (est.) | 14,250-42,750 | Unknown | Steroids, Traditional meds | Limited studies |
| **Australia** | 27,000,000 | Children | 3-5 | 5-10 | 162-270 | 0.1-0.2 | Observation, IVIG | High healthcare access |
| | | Adults | 2-4 | 12-20 | 432-864 | 0.1-0.3 | Rituximab, Steroids | Western patterns |
| **South Africa** | 62,000,000 | Children | 2-4 (est.) | 5-10 (est.) | 248-496 | Unknown | Observation, Steroids | Limited reporting infrastructure |
| | | Adults | 1-3 (est.) | 5-15 (est.) | 620-1,860 | Unknown | Steroids, Observation | Underreported chronic cases |
| **Russia** | 144,000,000 | Children | 2-4 (est.) | 5-10 (est.) | 576-1,152 | Unknown | Observation, Steroids | Limited specific data |
| | | Adults | 1-3 (est.) | 10-15 (est.) | 1,440-4,320 | Unknown | Steroids, IVIG | Extrapolated from EU trends |
| **Mexico** | 130,000,000 | Children | 3-5 (est.) | 5-10 (est.) | 780-1,300 | Unknown | Observation, IVIG | Latin American trends |
| | | Adults | 2-4 (est.) | 10-15 (est.) | 1,560-3,120 | Unknown | Steroids, Observation | Emerging data |
| **Italy** | 59,000,000 | Children | 3-5 | 5-10 | 354-590 | 0.1 | Observation, Steroids | EU registry data |
| | | Adults | 2-4 | 15-20 | 944-1,888 | 0.1-0.3 | Rituximab, Splenectomy | Higher adult prevalence |
| **Nigeria** | 230,000,000 | Children | 2-4 (est.) | 5-10 (est.) | 920-1,840 | Unknown | Observation, Steroids | Very limited data |
| | | Adults | 1-3 (est.) | 5-10 (est.) | 2,300-6,900 | Unknown | Steroids, Observation | Underreporting likely |
| **France** | 65,000,000 | Children | 3-5 | 5-10 | 390-650 | 0.1 | Observation, IVIG | Nationwide study (2009-2011) |
| | | Adults | 2-3 | 15-20 | 1,040-1,560 | 0.1-0.2 | Steroids, Rituximab | High adult chronicity |
| **Spain** | 47,000,000 | Children | 3-5 (est.) | 5-10 (est.) | 282- | | | |

| | | | | | | | | |
|---|---|---|---|---|---|---|---|---|
| | | | | | 470 | 0.1 | Observation, Steroids | EU trends applied |
| | | Adults | 2-4 (est.) | 12-20 (est.) | 752-1,504 | 0.1-0.3 | Rituximab, Splenectomy | Similar to Italy |
| **South Korea** | 51,000,000 | Children | 3-5 (est.) | 5-10 (est.) | 306-510 | 0.1 | Observation, IVIG | East Asian trends, H. pylori link |
| | | Adults | 2-4 (est.) | 15-20 (est.) | 816-1,632 | 0.1-0.2 | Steroids, Rituximab | Aging population impact |
| **Egypt** | 112,000,000 | Children | 2-4 (est.) | 5-10 (est.) | 448-896 | Unknown | Observation, Steroids | Limited data, regional estimate |
| | | Adults | 1-3 (est.) | 5-15 (est.) | 1,120-3,360 | Unknown | Steroids, Observation | Underreported |
| **Argentina** | 46,000,000 | Children | 3-5 (est.) | 5-10 (est.) | 276-460 | Unknown | Observation, IVIG | Latin American patterns |
| | | Adults | 2-4 (est.) | 10-15 (est.) | 736-1,472 | Unknown | Steroids, IVIG | Emerging data |
| **Pakistan** | 245,000,000 | Children | 2-4 (est.) | 5-10 (est.) | 980-1,960 | Unknown | Observation, Steroids | Sparse data, high population |
| | | Adults | 1-3 (est.) | 5-15 (est.) | 2,450-7,350 | Unknown | Steroids, Observation | Underreported |
| **Thailand** | 70,000,000 | Children | 2-4 (est.) | 5-10 (est.) | 280-560 | Unknown | Observation, Steroids | Southeast Asian estimate |
| | | Adults | 1-3 (est.) | 10-15 (est.) | 700-2,100 | Unknown | Steroids, IVIG | Limited studies |

##### Methodology and Sources:
1. **Population**: 2025 projections from UN, World Bank, or national statistics (approximate).
2. **Incidence/Prevalence**:
   - **Developed Countries (US, UK, Japan, Canada, Germany, Australia, France, Italy, Spain, South Korea)**: Based on studies like US claims (2-6/100,000), Nordic registries (1.8-2.8/100,000), French data (2.9/100,000), and Japanese studies (3-4/100,000). Children: 3-6/100,000; Adults: 10-25/100,000 prevalence due to chronicity.
   - **Developing Countries (India, Brazil, China, South Africa, Russia, Mexico, Nigeria, Egypt, Argentina, Pakistan, Thailand)**: Estimated from global ranges (1-5/100,000 incidence, 5-15/100,000 prevalence), adjusted for underreporting and healthcare access.
3. **Estimated Cases**: `(Population × Incidence Rate) / 100,000`, using ranges.
4. **Mortality**: 0.1-0.5/100,000 in countries with data; unavailable elsewhere due to rarity and lack of reporting.
5. **Treatments**: Reflect regional practices—observation/IVIG for children, steroids/TPO-RA/splenectomy for adults.

##### Notes on Data:
- **Robust Data**: US, UK, Japan, France, Germany, Italy from studies/registries.

- **Estimates**: Countries like Egypt, Pakistan, Thailand, Nigeria lack specific ITP studies; data inferred from regional/global trends.
- **Age Trends**: Children have higher incidence (acute cases), adults higher prevalence (chronic cases).

**Supplementary Information 2**
**GROK3 based research summary on molecules in clinical trials for ITP**
*Done on March 31, 2025 4:25 PM CST*

Here are some molecules currently in clinical trials:

#### 1. Rilzabrutinib
- **Type**: Oral Bruton Tyrosine Kinase (BTK) inhibitor
- **Purpose**: Blocks the BTK enzyme to reduce platelet destruction in ITP
- **Status**: Phase 3 trials
- **Details**: Early trials show promising results, with many patients achieving a sustained increase in platelet counts.

#### 2. Fostamatinib
- **Type**: Oral spleen tyrosine kinase (Syk) inhibitor
- **Purpose**: Reduces platelet destruction by targeting immune cell signaling
- **Status**: Approved for chronic ITP, with ongoing trials to expand its use
- **Details**: Studies are exploring its effectiveness in broader patient groups and in combination therapies.

#### 3. Avatrombopag
- **Type**: Thrombopoietin receptor agonist (TPO-RA)
- **Purpose**: Stimulates platelet production
- **Status**: In trials to optimize its use for ITP
- **Details**: Focus is on patients who don’t respond to other treatments.

#### 4. Efgartigimod
- **Type**: Neonatal Fc receptor (FcRn) inhibitor
- **Purpose**: Lowers levels of autoantibodies that destroy platelets
- **Status**: Phase 3 trials for ITP
- **Details**: An investigational drug showing potential in reducing immune-mediated platelet loss.

#### 5. Sutimlimab
- **Type**: Complement inhibitor (targets C1s)
- **Purpose**: Reduces complement-mediated platelet destruction
- **Status**: Phase 2 trials
- **Details**: Early results suggest it could be a new option for ITP treatment.

#### 6. Mezagitamab
- **Type**: Monoclonal antibody targeting CD38
- **Purpose**: Depletes plasma cells to reduce autoantibody production
- **Status**: Early-phase trials
- **Details**: A novel approach still in the initial stages of testing.

These examples showcase a variety of molecules—small molecules, biologics, and monoclonal antibodies—being tested in clinical trials. Clinical trials generally progress through phases: Phase 1 (safety), Phase 2 (efficacy and dosing), Phase 3 (large-scale effectiveness), and sometimes Phase 4 (post-approval studies). The molecules listed above are at different stages, and their success will depend on trial outcomes.

**Supplementary Information 3**
**GROK3 based research summary on clinical trials underway on ITP**
*Done on March 31, 2025 4:35 PM CST*
#### Key Points
- Research suggests several ongoing clinical trials for Immune Thrombocytopenia (ITP) in 2025, focusing on new treatments like ianalumab and mezagitamab.
- It seems likely that these trials aim to improve platelet counts and manage symptoms, with some showing promising results.
- The evidence leans toward increased focus on novel agents targeting various disease mechanisms, potentially offering new options for patients.

#### Overview
Immune Thrombocytopenia (ITP) is a condition where the immune system attacks platelets, leading to low platelet counts and increased bleeding risk. Clinical trials in 2025 are exploring new treatments to address this, offering hope for better management.

#### Ongoing Trials
Several trials are currently underway, including:
- **VAYHIT1 and VAYHIT2** by Novartis, testing ianalumab with corticosteroids or eltrombopag for adults with ITP.
- A HUTCHMED trial evaluating HMPL-523, a selective Syk inhibitor, for adult ITP patients.
- Takeda's mezagitamab, with positive phase 2b results and a planned phase 3 trial, showing improved platelet responses.

#### Unexpected Detail: Pediatric Focus
An unexpected finding is the inclusion of pediatric trials, such as one comparing eltrombopag to standard front-line management for newly diagnosed ITP in children, highlighting efforts to address younger patients.

For more details, visit [clinicaltrials.gov](https://clinicaltrials.gov/) for trial specifics.

---

#### Survey Note: Comprehensive Update on ITP Clinical Trials as of March 31, 2025

This note provides a detailed overview of the latest developments in clinical trials for Immune Thrombocytopenia (ITP), a rare autoimmune disorder characterized by low platelet counts and increased bleeding risk. The information is compiled from various sources, including academic articles, clinical trial registries, and pharmaceutical company announcements, ensuring a comprehensive and up-to-date analysis as of the current date.

##### Background on ITP and Clinical Trials
ITP, or Immune Thrombocytopenia, results from the immune system mistakenly attacking platelets, leading to thrombocytopenia and potential bleeding complications. Historically, treatments have included corticosteroids and splenectomy, but recent decades have seen the introduction of thrombopoietin receptor agonists, revolutionizing the treatment landscape. Despite these advances, significant unmet needs remain, prompting ongoing research into novel and repurposed agents. Clinical trials in 2025 are focusing on early-phase interventions, aiming to modify the disease course and target previously unaddressed pathophysiological mechanisms, such as platelet autoantibody recycling, B-cell maturation, long-lived plasma cells, and the complement system.

##### Detailed Trial Updates
The following table summarizes key ongoing and recently updated ITP clinical trials, based on available data:

| **Trial Name/Code** | **Phase** | **Target Population** | **Treatment** | **Purpose** | **Status** | **URL** |
| --- | --- | --- | --- | --- | --- | --- |
| VAYHIT1 (NCT05653349) | Phase 3 | Adults (≥ 18 years) with newly diagnosed primary ITP within last 3 months, platelet count < 30G/L before corticosteroids, ≥ 50G/L on corticosteroids prior to

randomization | Ianalumab (higher or lower dose) vs placebo, with first-line corticosteroids | Assess if ianalumab maintains platelet count ≥ 30G/L without rescue treatment or new ITP therapy, prolongs time to treatment failure | Ongoing | [clinicaltrials.gov](https://clinicaltrials.gov/study/NCT05653349) |
| HUTCHMED (NCT06291415) | Not specified | Adults diagnosed with ITP ≥ 3 months ago, intolerance/insufficient response/recurrence after ≥ 1 anti-ITP standard drug, platelet count < 30,000 μL for ≥ 2 visits during screening, no count > 35,000 μL in 2 weeks before enrollment | HMPL-523 tablets once daily for 24 weeks, across dose levels | Evaluate safety, tolerability, and efficacy of HMPL-523 (selective Syk inhibitor) in ITP | Ongoing | [clinicaltrials.gov](https://clinicaltrials.gov/study/NCT06291415) |
| VAYHIT2 (NCT05653219) | Phase 3 | Patients with primary ITP requiring second-line therapy with TPO-RA, eligible for eltrombopag | Ianalumab (higher or lower dose) vs placebo, with eltrombopag | Assess if ianalumab with eltrombopag improves long-term outcome by prolonging time to treatment failure compared to eltrombopag monotherapy | Ongoing | [clinicaltrials.gov](https://clinicaltrials.gov/study/NCT05653219) |
| Eltrombopag vs Standard Front Line Management | Phase 3 | Children (ages 1 to <18 years) with newly diagnosed ITP | Eltrombopag vs standard front-line management | Compare efficacy and safety in newly diagnosed pediatric ITP | In progress, not accepting new patients | [UCSF trials](https://clinicaltrials.ucsf.edu/trial/NCT03939637) |
| Rilzabrutinib in Adults and Adolescents | Not specified | Adults and adolescents with persistent or chronic ITP, average platelet count <30,000/μL | Rilzabrutinib or placebo 400mg twice daily | Evaluate safety and efficacy in persistent or chronic ITP | In progress, not accepting new patients | [UCSF trials](https://clinicaltrials.ucsf.edu/trial/NCT04562766) |
| Mezagitamab (TAK-079) | Phase 2b (with planned Phase 3) | Adults with persistent or chronic primary ITP | Mezagitamab (100 mg, 300 mg, 600 mg) vs placebo, subcutaneous, once weekly for 8 weeks | Assess safety, tolerability, and efficacy, with focus on platelet response | Positive phase 2b results, phase 3 planned for FY2024 | [Takeda news](https://www.takeda.com/newsroom/newsreleases/2024/late-breaking-data-from-phase-2b-study-of-mezagitamab/) |

##### Analysis of Trial Focus and Innovations
The trials listed reflect a shift from historical approaches, which focused on indefinite treatment for persistent or chronic ITP, to a broader spectrum including early-phase interventions. Notably, ianalumab trials (VAYHIT1 and VAYHIT2) by Novartis target B-cell maturation and differentiation, aiming to prolong time to treatment failure, particularly in combination with established therapies like eltrombopag. The HUTCHMED trial with HMPL-523 introduces a selective Syk inhibitor, targeting a novel mechanism to enhance platelet production and reduce immune-mediated destruction.

An unexpected detail is the inclusion of pediatric-focused trials, such as the eltrombopag comparison study, which addresses a critical gap in treatment options for younger patients. This is significant given the challenges of managing ITP in children, where long-term outcomes and quality of life are key concerns.

Mezagitamab, developed by Takeda, stands out with its phase 2b results showing rapid and sustained platelet count increases above the 50,000/μL threshold, with over 81% of patients on the 600mg dose achieving a complete platelet response. This drug, a fully human IgG1 monoclonal antibody targeting CD38-expressing cells, has a favorable safety profile, with no new safety signals, and is poised for a global phase 3 trial, potentially transforming ITP treatment.

##### Additional Insights from Literature
Recent literature, such as the 2025 update on ITP clinical trials published in August 2024 ([PubMed](https://pubmed.ncbi.nlm.nih.gov/39105413/)), highlights the ongoing development of investigational agents targeting platelet autoantibody recycling, long-lived plasma cells, and the complement system. This aligns with the trial data, indicating a trend toward personalized and mechanism-based therapies. The UCSF trials page, updated in February 2025, confirms the ongoing nature of rilzabrutinib and pediatric eltrombopag studies, emphasizing multicenter efforts to broaden the evidence base.

##### Implications for Patients and Future Directions
These trials suggest a promising future for ITP management, with potential new treatments that could offer sustained platelet responses and reduced reliance on corticosteroids. Patients should discuss

eligibility with healthcare providers, considering the global reach of these trials, as seen in the VAYHIT studies and Takeda's planned phase 3 for mezagitamab. The focus on both adult and pediatric populations, as well as novel mechanisms, indicates a comprehensive approach to addressing ITP's challenges.

This survey note encapsulates all available details from recent sources, ensuring a thorough understanding of the current ITP clinical trial landscape as of March 31, 2025.

##### Key Citations
- [2025 update on clinical trials in immune thrombocytopenia PubMed article](https://pubmed.ncbi.nlm.nih.gov/39105413/)
- [UCSF Immune Thrombocytopenia Clinical Trials for 2025 page](https://clinicaltrials.ucsf.edu/immune-thrombocytopenia)
- [2025 update on clinical trials in immune thrombocytopenia Wiley article](https://onlinelibrary.wiley.com/doi/full/10.1002/ajh.27448)
- [VAYHIT1 clinical trial on clinicaltrials.gov](https://clinicaltrials.gov/study/NCT05653349)
- [HUTCHMED clinical trial on clinicaltrials.gov](https://clinicaltrials.gov/study/NCT06291415)
- [VAYHIT2 clinical trial on clinicaltrials.gov](https://clinicaltrials.gov/study/NCT05653219)
- [Eltrombopag vs standard front-line management trial on UCSF](https://clinicaltrials.ucsf.edu/trial/NCT03939637)
- [Rilzabrutinib trial on UCSF](https://clinicaltrials.ucsf.edu/trial/NCT04562766)
- [Takeda Mezagitamab phase 2b results news release](https://www.takeda.com/newsroom/newsreleases/2024/late-breaking-data-from-phase-2b-study-of-mezagitamab/)
